# Quantitative secretome analysis establishes the cell type-resolved mouse brain secretome

**DOI:** 10.1101/2020.05.22.110023

**Authors:** Johanna Tüshaus, Stephan A. Müller, Evans Sioma Kataka, Jan Zaucha, Laura Sebastian Monasor, Minhui Su, Gökhan Güner, Georg Jocher, Sabina Tahirovic, Dmitrij Frishman, Mikael Simons, Stefan F. Lichtenthaler

## Abstract

To understand how cells communicate in the nervous system, it is essential to define their secretome, which is challenging for primary cells because of large cell numbers being required. Here, we miniaturized secretome analysis by developing the high-performance secretome-protein-enrichment-with-click-sugars method (hiSPECS). To demonstrate its broad utility, hiSPECS was used to identify the secretory response of brain slices upon LPS-induced neuroinflammation and to establish the cell type-resolved mouse brain secretome resource using primary astrocytes, microglia, neurons and oligodendrocytes. This resource allowed mapping the cellular origin of CSF proteins and revealed that an unexpectedly high number of secreted proteins *in vitro* and *in vivo* are proteolytically-cleaved membrane protein ectodomains. Two examples are neuronally secreted ADAM22 and CD200, which we identified as substrates of the Alzheimer-linked protease BACE1. hiSPECS and the brain secretome resource can be widely exploited to systematically study protein secretion, brain function and to identify cell type-specific biomarkers for CNS diseases.

## Introduction

Protein secretion is essential for inter-cellular communication and tissue homeostasis of multicellular organisms and has a central role in development, function, maintenance and inflammation of the nervous system. Proteins secreted from cells are referred to as the secretome and comprise secreted soluble proteins, such as insulin, granulins, APOE and extracellular matrix proteins (e.g. neurocan, fibronectin). The secretome also comprises the extracellular domains of membrane proteins, e.g. growth factors, cytokines, receptors and cell adhesion proteins (e.g. neuregulin, NCAM, N-cadherin), which are proteolytically generated by mostly membrane-bound proteases and secreted in a cellular process called ectodomain shedding (Lichtenthaler, Lemberg et al., 2018). However, it is largely unknown to what extent ectodomain shedding contributes to total protein secretion and how this differs between cell types in the brain.

Omics’ approaches have generated large collections of mRNA and protein abundance data across the different cell types of the brain, e.g.(Sharma, Schmitt et al., 2015, Zhang, Chen et al., 2014). In contrast, little is known about the proteins that are secreted from brain cells and whether – in parallel to their broad expression in different brain cell types – they are secreted from multiple brain cell types or instead are secreted in a cell type-specific manner *in vitro, ex vivo* (e.g. organotypic slice culture), *and in vivo*. Cerebrospinal fluid (CSF) constitutes an *in vivo* brain secretome and is an easily accessible body fluid widely used for studying brain (patho-)physiology and measuring and identifying disease biomarkers (Johnson, Dammer et al., 2020, Olsson, Lautner et al., 2016, Zetterberg & Bendlin, 2020), but it is largely unknown which cell type the CSF proteins are secreted from, because no systematic brain cell type-specific protein secretion studies are available.

Dysregulated protein secretion and shedding is linked to neurologic and psychiatric diseases, including neurodegeneration, e.g. APP, APOE, SORL1 and TREM2 in Alzheimeŕs disease (AD), or the prion protein (PRNP) in Prion disease (Lichtenthaler et al., 2018). Thus, identification and quantification of secretomes does not only allow understanding of biological processes under physiological conditions, but also contributes to unravelling the molecular basis of diseases and identification of drug targets and biomarkers, such as shed TREM2 for AD (Ewers, Franzmeier et al., 2019, Schindler, Li et al., 2019, Suarez-Calvet, Araque Caballero et al., 2016).

Systematic identification and quantification of secretome proteins is commonly done using conditioned medium of a (primary) cell type and its analysis by mass spectrometry-based proteomics. A major challenge is the low concentration of secreted proteins within the conditioned medium(Schira-Heinen, Grube et al., 2019). Therefore, many studies concentrate medium from tens of millions of cells (Kleifeld, Doucet et al., 2010, Kleifeld, Doucet et al., 2011, Kuhn, Koroniak et al., 2012, Schlage, Kockmann et al., 2015, Wiita, Seaman et al., 2014). However, such numbers are often not available for primary cells, such as microglia, where on average one million cells may be purified from the brain of individual adult mice. Thus, miniaturized secretome analysis methods are required. A second major challenge is the large dynamic range of secretomes, in particular when cells are cultured in the presence of serum or serum-like supplements, which are highly abundant in proteins (most notably albumin), hampering the detection of endogenous, cell-derived secreted proteins, whose protein levels are typically orders of magnitude lower(Eichelbaum, Winter et al., 2012). Therefore, cells are often cultured under serum- and protein-free starvation conditions(Deshmukh, Cox et al., 2015, Kleifeld et al., 2010, Meissner, Scheltema et al., 2013), which, however, strongly alters secretome composition and may induce cell-death(Eichelbaum et al., 2012). An alternative approach, compatible with cell culture in the presence of serum or serum-like supplements, is to metabolically label the cell-derived, but not the exogenous serum proteins with analogs of methionine or sugars that are incorporated into the protein backbone or glycan structures, of newly synthesized cellular proteins, respectively (Eichelbaum et al., 2012, Kuhn et al., 2012). However, even these established methods still require extensive secretome fractionation, either at the protein or peptide level (Eichelbaum et al., 2012, Kuhn et al., 2012) and, thus result in laborious sample preparation, extensive mass spectrometry measurement times and the requirement of large amounts of samples, which may not be available from primary cells or tissues.

Here, we developed the ‘high-performance secretome protein enrichment with click sugars’ (hiSPECS) method, which down-scales and speeds up secretome analysis and now allows secretome analysis of primary brain cells from single mice. We applied hiSPECS to determine the cell type-resolved mouse brain secretome, which establishes a resource for systematically studying protein secretion and shedding in the brain. Broad applicability of hiSPECS and the resource is demonstrated a) by gaining new insights into the extent of cell type-specific protein secretion and shedding, b) by identifying new substrates for the protease BACE1, a major drug target in AD, c) by determining the cellular origin of proteins in CSF and d) by revealing that LPS-induced inflammatory conditions in organotypic brain slices do not only lead to inflammatory protein secretion from microglia, but instead induce to a systemic secretory response from multiple cell types in brain slices.

## Results

### Development of hiSPECS and benchmarking against SPECS

To enable secretome analysis of primary brain cell types from single mice under physiological conditions (i.e. in the presence of serum-like supplements), we miniaturized the previously established SPECS method, which required 40 million cells per experiment (Kuhn et al., 2012). We introduced four major changes (Fig. 1A, see Supplementary Fig. 1A for a detailed comparison of hiSPECS versus SPECS). First, after labeling of cells with N-azido-mannosamine (ManNAz), an azido group-bearing sugar, secretome glycoproteins were enriched from the conditioned medium with lectin-based precipitation using concanavalin A (ConA). This strongly reduced albumin, which is not a glycoprotein (Supplementary Fig. 1B). Because the majority of soluble secreted proteins and most of the membrane proteins – which contribute to the secretome through shedding – are glycosylated (Kuhn, Colombo et al., 2016), hiSPECS can identify the major fraction of all secreted proteins. Second, we selectively captured the azido-glycoproteins by covalent binding to magnetic dibenzylcyclooctyne (DBCO)-alkyne-beads using copper-free click chemistry. This allowed stringent washing to reduce contaminating proteins. Third, on-bead digestion of the captured glycoproteins was performed to release tryptic peptides for mass spectrometry-based label-free protein quantification (LFQ). Fourth, mass spectrometry measurements were done on a Q-Exactive HF mass spectrometer using either data-dependent acquisition (DDA) or the more recently developed data-independent acquisition (DIA)(Bruderer, Bernhardt et al., 2015, Gillet, Navarro et al., 2012, Ludwig, Gillet et al., 2018)(Supplementary Fig.1C).

**Fig. 1:**
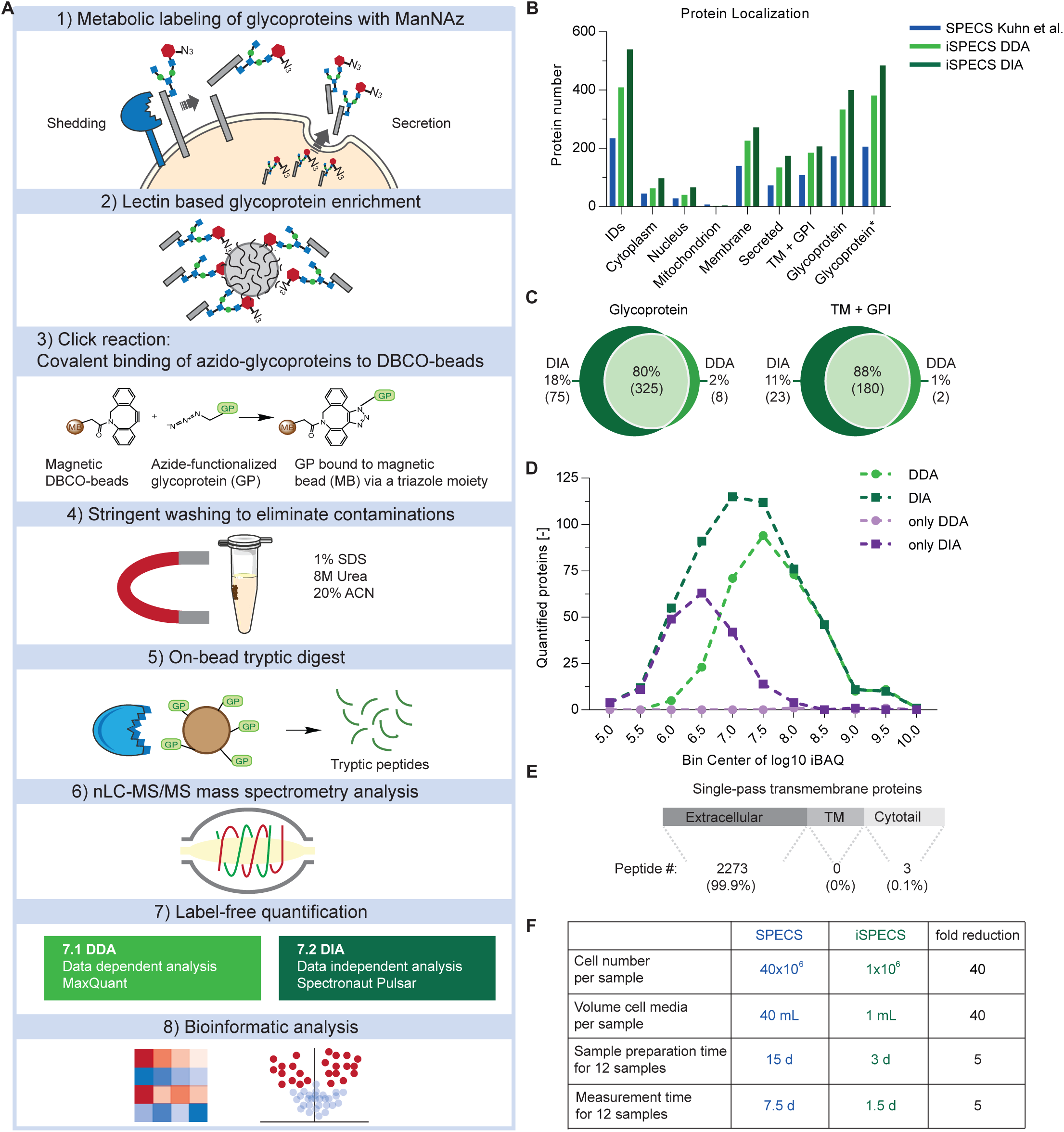
Workflow of the hiSPECS method and benchmarking against SPECS. **A)** Graphical illustration of the hiSPECS workflow. Cells are metabolically labeled with N-azido-mannosamine (ManNAz), an azido group-bearing sugar, which is metabolized in cells and incorporated as azido-sialic acid into N- and O-linked glycans of newly synthesized glycoproteins, but not into exogenously added serum proteins. **B)** Bar chart indicating protein quantification, protein localization (according to UniProt) in the secretome of primary neurons, comparing the previous SPECS (blue) to the new hiSPECS method using DDA (light green) or DIA (dark green). Proteins were counted if quantified in at least 9 of the 11 biological replicates of hiSPECS or 4 of 5 biological replicates of the previous SPECS paper (Kuhn et al., 2012). The category glycoprotein* includes UniProt annotations and proteins annotated as glycoproteins in previous papers(Fang et al., 2016, Joshi et al., 2018, Liu et al., 2017, Zielinska et al., 2010). Proteins can have multiple UniProt annotations, e.g. for APP membrane, TM, cytoplasm and nucleus, because distinct proteolytic fragments are found in different organelles, so that some proteins in categories cytoplasm and nucleus may overlap with the categories secreted and TM+GPI. **C)** Venn diagram comparing the number of protein groups quantified (5/6 biological replicates) by DDA versus DIA of the same samples. Left panel: glycoprotein. Right panel: TM and GPI proteins, which are potentially shed proteins. **D)** Distribution of quantified proteins with DDA and DIA (at least in 3 of 6 biological replicates). The number of quantified proteins is plotted against the log10 transformed label-free quantification (LFQ) values with a bin size of 0.5. The number of quantified proteins per bin for DDA and DIA are indicated in light and dark green, respectively. Proteins that were only quantified with DDA or DIA are colored in light and dark purple, respectively. DIA provides additional quantifications for low abundant proteins and extends the dynamic range for protein quantification by almost one order of magnitude. **E)** All tryptic peptides identified from TM proteins were mapped onto their protein domains. Only 0.1% of the peptides mapped to intracellular domains. Demonstrating that secretome proteins annotated as TM proteins comprise the shed ectodomains, but not the full-length forms of the proteins. **F)** Comparison of SPECS and hiSPECS method with regard to cell number, volume of culture media, sample preparation time and mass spectrometer measurement time. TM: single-pass transmembrane protein; GPI: Glycosylphosphatidylinositol-anchored membrane protein.

To benchmark hiSPECS against the previous SPECS protocol, we collected the secretome of primary murine neurons in the presence of a serum supplement as before (Kuhn et al., 2012), but used only one million neurons (40-fold fewer cells compared to SPECS). Despite this miniaturization, hiSPECS quantified on average 186% and 236% glycoproteins in DDA and DIA mode, respectively, compared to the previous SPECS dataset (Kuhn et al., 2012) (according to UniProt and four previous glycoproteomic studies; Fig. 1B; Supplementary Fig.1D,E; Supplementary Table 1) (Fang, Wang et al., 2016, Joshi, Jorgensen et al., 2018, Liu, Zeng et al., 2017, Zielinska, Gnad et al., 2010). Furthermore, the DIA method provided 18% more quantified glycoproteins and 11% shed transmembrane proteins compared to DDA (Fig. 1C). Due to the superiority of DIA over DDA in secretome analysis we focused on hiSPECS DIA throughout the manuscript (Fig. 1D). 99.9% (2273/2276) of unique tryptic peptides identified from single-pass transmembrane proteins mapped to their known or predicted protein ectodomains. This is an important quality control which demonstrates that the secretome contains proteolytically shed transmembrane ectodomains and not full-length transmembrane proteins and that hiSPECS reliably identifies secretome-specific proteins (Fig. 1E).

The novel hiSPECS procedure does not require further protein or peptide fractionation. Thus, sample preparation time and mass spectrometer measurement time were both reduced 5-fold compared to previously required times (Fig. 1F). Importantly, hiSPECS also improved the reproducibility of protein LFQ among different biological replicates, which is reflected by an average Pearson correlation coefficient of 0.97 using hiSPECS in comparison to 0.84 with the previous procedure (Supplementary Fig.1F). Taken together, hiSPECS outperforms SPECS with regard to the required number of cells, sensitivity, reproducibility, protein coverage, sample preparation and mass spectrometry measurement time.

### Cell type-resolved mouse brain secretome resource

Secreted soluble and shed proteins have key roles in signal transduction, but their cellular origin is often unclear as they may be expressed by multiple cell types. Thus, we used hiSPECS to establish a resource of the brain secretome in a cell type-resolved manner, focusing on the four major brain cell types – astrocytes, microglia, neurons and oligodendrocytes (Fig. 2A; Supplementary Table 2). One million primary cells of each cell type were prepared from individual mouse brains. For further analysis, we focused on proteins detected in the secretome of at least five out of six biological replicates of at least one cell type. This yielded 995 protein groups in the secretome, with microglia having on average the largest (753) and astrocytes the smallest (503) number of protein groups (Fig. 2B). GO cellular compartment analysis revealed extracellular region to be the most enriched term underlining the quality of our secretome library (Supplementary Fig. 2A,B). An additional quality control measure was the enrichment of known cell type-specifically secreted marker proteins such as LINGO1, SEZ6 and L1CAM in the neuronal secretome (Supplementary Fig. 2C). Importantly, the secretome analysis identified 111 proteins (Fig. 2C) that were not detected in a previous proteomic study by Sharma *et al*. (Sharma et al., 2015) (Supplementary Table 3), that identified >10,000 proteins in the lysates of the same primary mouse brain cell types, which had been taken in culture as in our study (Supplementary Table 2). This includes soluble proteins (e.g. CPN1, TIMP1) and shed ectodomains (e.g. ADAM19, CRIM1, FRAS1) and even the protein GALNT18, which was assumed to be a pseudogene that is not expressed as a protein. Together, this demonstrates that secretome analysis is complementary to lysate proteomics in order to identify the whole proteome expressed in an organ.

**Fig. 2:**
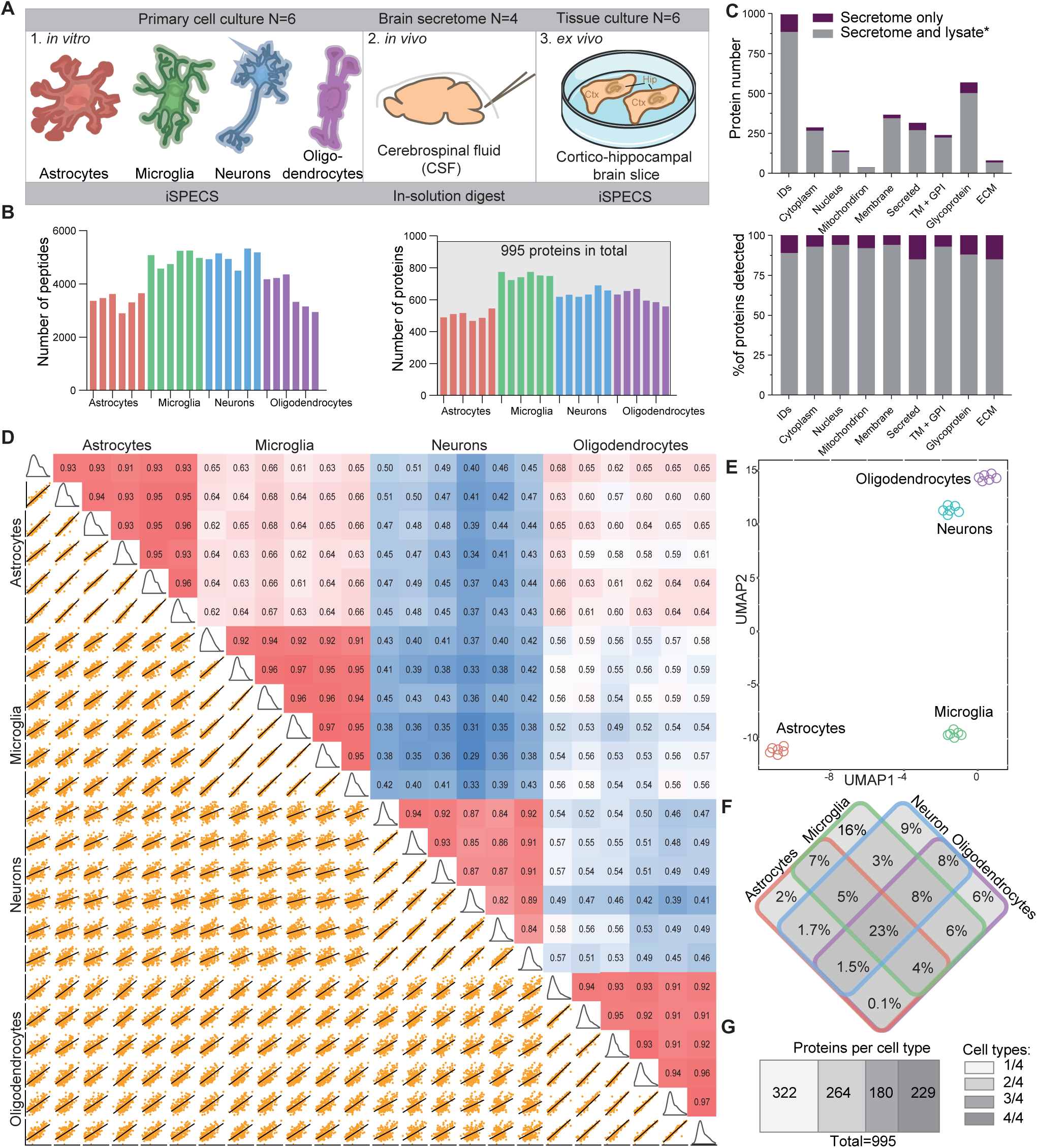
Cell type-resolved mouse brain secretome resource. **A)** Illustration of the proteomic resource including secretome analysis of primary murine astrocytes, microglia, neurons, oligodendrocytes, cerebrospinal fluid (CSF) analysis and brain slices (hippocampus (Hip), cortex (Ctx)). **B)** Number of peptides (left) and protein groups (right) quantified in the secretome of the investigated brain cell types with the hiSPECS DIA method. Proteins quantified in at least 5 out of 6 biological replicates in at least one cell type are considered. In total 995 protein groups were detected. **C)** 995 proteins were quantified and identified (ID) in the hiSPECS secretome resource. The grey part of each column indicates how many proteins were also detected in the lysates of the same four cell types as analyzed in a previous proteome dataset(Sharma et al., 2015). This reveals that 111 proteins (purple) were only detected in the secretome. Relative to all the secretome proteins covered by the lysate study (grey) the most enriched UniProt annotation of the newly identified proteins (purple) was secreted and extracellular matrix (15%) (lower panel). **D)** Correlation matrix showing the relationship between the different brain cell types. The matrix shows the Pearson correlation coefficient (red indicates a higher, blue a lower correlation) and the correlation plots of the log2 LFQ intensities of the secretome of astrocytes, neurons, microglia and oligodendrocytes processed with the hiSPECS method. **E)** UMAP (Uniform Manifold Approximation and Projection) plot showing the segregation of the brain cell type secretomes based on LFQ intensities of quantified proteins. **F)** Venn diagram illustrating the percentage of the secretome proteins which were secreted from only one or multiple cell types. Proteins quantified in at least 5 biological replicates of one cell type were considered. **G)** Bar graph indicating the number of protein groups, which were detected in one, two, three or all cell types with at least 5 biological replicates.

Based on LFQ intensity values, the Pearson correlation coefficients between the six biological replicates of each cell type showed on average an excellent reproducibility with a value of 0.92 (Fig. 2D). In strong contrast, the correlation between different cell types was dramatically lower (0.29 - 0.68), indicating prominent differences between the cell type-specific secretomes (Fig. 2D-F Supplementary Fig. 2D,3). In fact, about one third (322/995) of the secretome proteins were consistently detected (5/6 biological replicates) in the secretome of only one cell type and in fewer replicates or not at all within the other cell type secretomes (Fig. 2G), highlighting the unique cell type-characteristic secretome fingerprint of each cell type (Supplementary Table 4).

Cell type-specific protein secretion was visualized in a heat map (Fig. 3A, Supplementary Fig. 4A). Gene ontology analysis of the enriched secretome proteins revealed functional clusters corresponding to the known functions of the four individual cell types. For example, metabolic process, gliogenesis and immune response was preferentially secreted from astrocytes (e.g. IGHM, CD14, LBP). Functional clusters autophagy and phagocytosis were secreted from microglia (e.g. TGFB1, MSTN, TREM2), in agreement with key microglial functions. Neuron-preferentially secreted proteins belonged to the neuron-specific categories axon guidance, trans-synaptic signaling and neurogenesis (e.g. NCAN, CHL1, SEZ6). Oligodendrocyte-specifically secreted proteins (e.g. OMG, ATRN, TNFRSF21) fell into categories lipid metabolic process and myelination, in agreement with the role of oligodendrocytes in myelin sheet formation. This demonstrates that cell function can be obtained by a celĺs secretome (Fig. 3A, Supplementary Fig.4B).

**Fig. 3:**
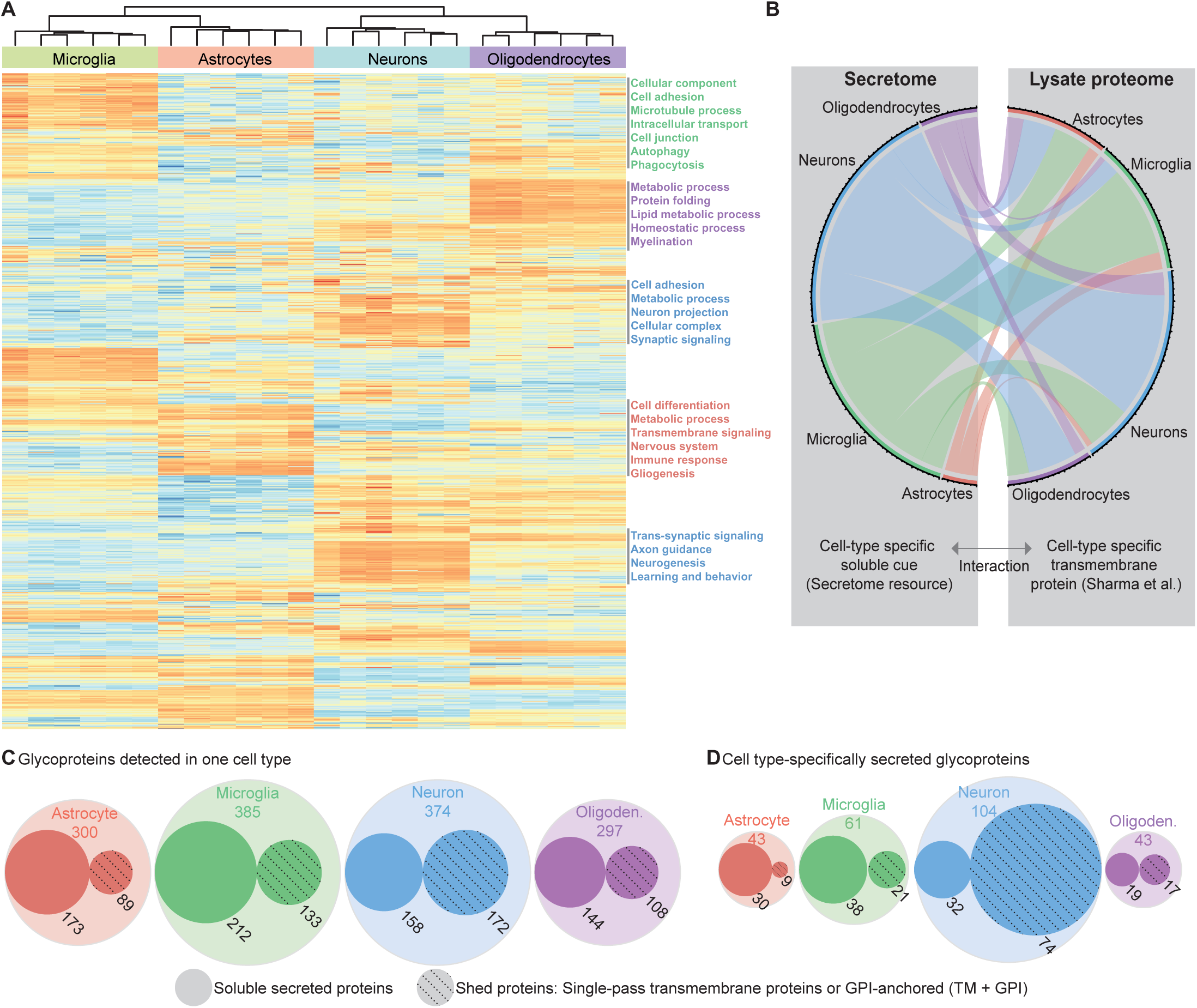
Cell type-specific enrichment of proteins in the secretome of brain cells. **A)** Heat map of the hiSPECS library proteins across the four cell types from hierarchical clustering. For missing protein data, imputation was performed. Rows represent the 995 proteins and columns represent the cell types with their replicates. The colors follow the z-scores (blue low, white intermediate, red high). Functional annotation clustering with DAVID 6.8(Huang da et al., 2009a, Huang da et al., 2009b) for the gene ontology category biological process (FAT) of protein clusters enriched in one cell type are indicated on the right sorted by the enrichment score (background: all proteins detected in the hiSPECS brain secretome resource). **B)** Interaction map of the hiSPECS secretome and the published lysate proteome data(Sharma et al., 2015) illustrating the cellular communication network between the major brain cell types. Interaction pairs are based on binary interaction data downloaded from UniProt and BioGRID databases (Chatr-Aryamontri et al., 2015, UniProt Consortium, 2018). Cell type-specifically secreted proteins of the hiSPECS secretome resource were mapped to their interaction partners if the interaction partners also show cell type-specificity in lysates of one brain cell type in the proteome data(Sharma et al., 2015) (2.5-fold pairwise) and are annotated as transmembrane protein in UniProt (Supplementary Table 5). **C)** Visualization of the glycoproteins detected in at least 5 of 6 biological replicates of the four brain cell types. Soluble secreted or potentially shed proteins are indicated for each cell type. The radius of the circles resembles the protein count. Shed proteins: includes proteins annotated as single-pass transmembrane and glycosylphosphatidylinositol (GPI)-anchored proteins, of which the shed ectodomain was found in the secretome; Sec: soluble secreted. **D)** Visualization of the cell type specific (CTS) glycoproteins as in C), either specifically secreted by one cell type in at least 5 biological replicates and no more than 2 biological replicates in another cell type or 5-fold enriched according to pairwise comparisons to the other cell types (Supplementary Table 4).

Secreted proteins may act as soluble cues to signal to other cells. To unravel the inter-cellular communication between secreted proteins and transmembrane proteins acting as potential binding partners/receptors, we mapped known interaction partners (from UniProt and BioGRID (Chatr-Aryamontri, Breitkreutz et al., 2015, UniProt Consortium, 2018)), but now in a cell type-resolved manner (Fig. 3B and Supplementary Table 5). Besides known cell type-specific interactions, such as between neuronally secreted CD200 and its microglia-expressed receptor CD200R1 (Yi, Zhang et al., 2016), we also detected new cell type-specific interactions, e.g. between ADIPOQ (adiponectin) and CDH13. Adiponectin is a soluble anti-inflammatory adipokine with key functions in metabolism, but also in neurogenesis and neurodegeneration (Lee, Cheng et al., 2019). Although adiponectin is thought to be secreted exclusively from adipocytes and assumed to reach the brain by crossing the blood-brain-barrier, our resource and the cell type-resolved interaction map reveal that adiponectin can also be produced and secreted from brain cells (oligodendrocytes). The binding to one of its receptors, cadherin13 (neurons), establishes a new interaction between oligodendrocytes and neurons, which may have an important role in controlling the multiple, but not yet well understood, adiponectin functions in the brain.

Besides pronounced cell type-specific protein secretion, another key insight obtained from our secretome resource is that ectodomain shedding of membrane proteins strongly contributes to the composition of the secretome (43%, 242/568 glycoproteins according to UniProt). The extent differed between the major mouse brain cell types (Fig. 3C) and became even more evident when focusing on the cell type-specifically secreted proteins. More than two thirds (71%, 74/104) of the neuron-specifically secreted proteins were shed ectodomains (Fig. 3D), i.e. proteins annotated as transmembrane or GPI-anchored proteins, of which we almost only (99.9%) identified peptides from the ectodomain (Fig. 1E). The neuronally shed ectodomains contain numerous trans-synaptic signaling and cell adhesion proteins (NRXN1, SEZ6, CNTNAP4) indicating that shedding is an important mechanism for controlling signaling and synaptic connectivity in the nervous system. In contrast to neurons, shedding appeared quantitatively less important in astrocytes, where only 21% (9/43) of the cell type-specifically secreted proteins were shed ectodomains (Fig. 3D). In contrast, soluble secreted proteins are particularly relevant for astrocytes and microglia where they constituted 70% (30/43) and 62% (38/61) of the secretome proteins, respectively, and contain numerous soluble proteins with functions in inflammation, e.g. complement proteins and TIMP1 (astrocytes) and GRN, MMP9, PLAU, TGFB1 (microglia). Importantly, several soluble secreted proteins were expressed at similar levels in different brain cell types, but predominantly secreted from only one, suggesting that cell type-specific protein secretion may be an important mechanism to control brain inflammation. Examples are the astrocyte-secreted complement factor B and the microglia-secreted LTBP4 (Supplementary Table 4).

### Mechanisms of cell type-specific protein secretion

Abundance levels in the lysate of the majority of proteins are similar among astrocytes, microglia, neurons and oligodendrocytes, with only around 15% of the proteins present in a cell type-specific manner (Sharma et al., 2015). In clear contrast, we observed that in the secretome of the same cell types, nearly half (420/995 proteins) of the secreted proteins were secreted in a cell type-specific manner (>5-fold enriched in one secretome compared to all three others or consistently detected in the secretome of only one cell type) (Supplementary Table 4), suggesting that cell type-specific protein secretion strongly contributes to functional differences between the four brain cell types.

An obvious mechanism explaining cell type-specific protein secretion is cell type-specific expression of the secreted soluble or shed proteins. Unexpectedly, however, a correlation with cell type-specific protein abundance in the same cell types (Sharma et al., 2015) was observed for only 20-30% of the cell type-specifically secreted proteins (Fig. 4A), including the TGF*β* coreceptor CD109 from astrocytes, the inflammatory proteins GRN, BIN2, TGF*β*1 from microglia, the cell adhesion proteins CD200, L1CAM, and the bioactive peptide secretogranin (CHGB) from neurons and CSPG4, PDGFR*α*, OMG and BCHE from oligodendrocytes (examples shown in Fig. 4A,B and Supplementary Fig.5). This demonstrates that cell type-specific expression is only a minor or only one of several mechanisms controlling cell type-specific protein secretion and shedding. In fact, the vast majority of cell type-specifically secreted proteins were equally expressed in two or more cell types (Fig. 4A).

**Fig. 4:**
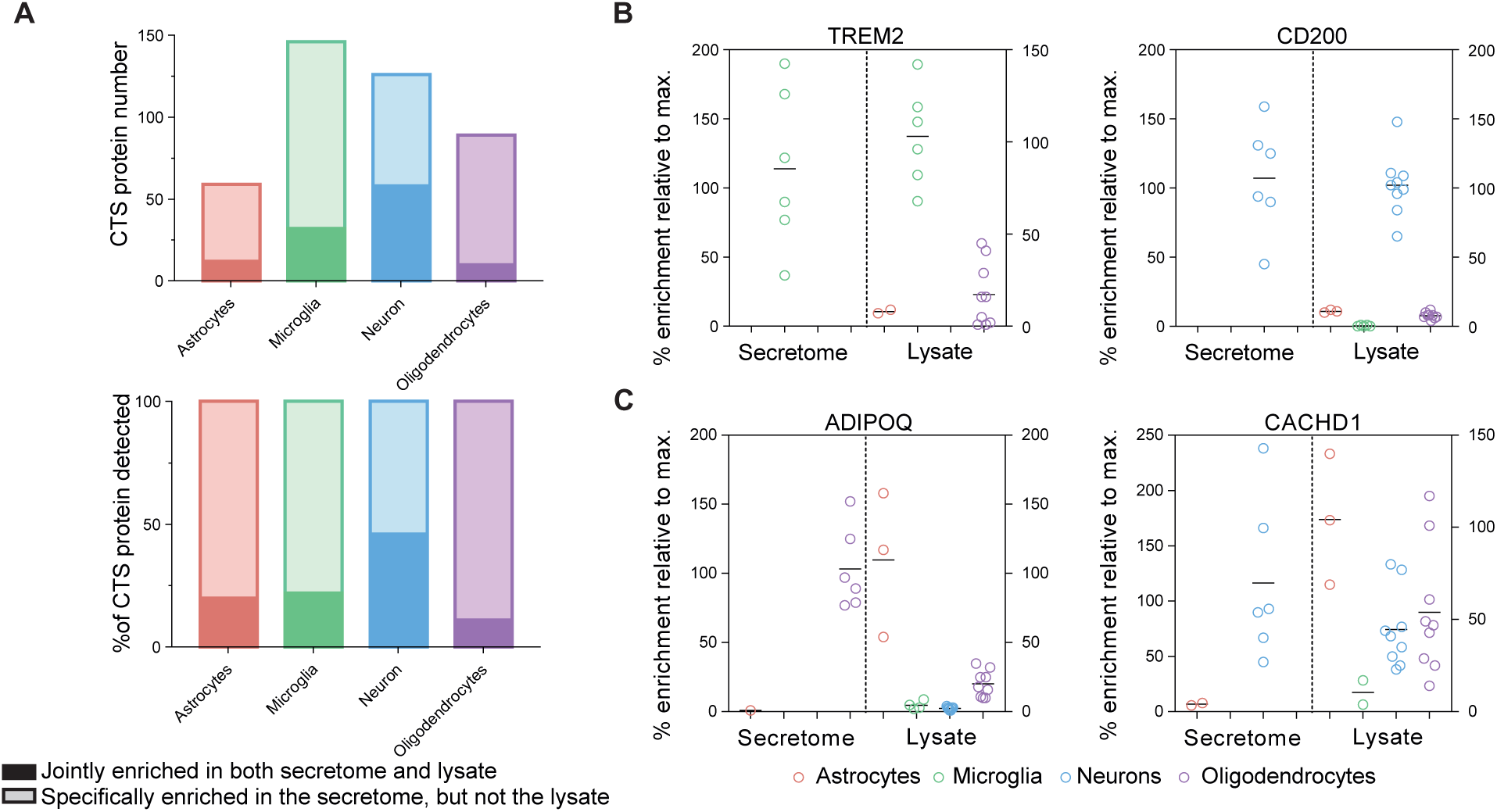
Protein levels in the brain cell secretome vs. lysate proteome. **A)** Bar graph of proteins specifically secreted from the indicated cell types. The solid part of the box indicates which fraction of proteins were predominantly enriched both in the secretome and the lysate of the indicated cell type, suggesting that cell type-specific secretion results from cell type-specific protein synthesis. The light part of the box shows the fraction of proteins that were specifically secreted from the indicated cell type, although this protein had similar levels in the lysate of multiple cell types, indicating cell type-specific mechanisms of secretion or shedding. Lysate protein levels were extracted from (Sharma et al., 2015). **B-C)** Comparison of the hiSPECS secretome resource and lysate data by (Sharma et al., 2015). The % enrichment is indicated normalized to the average of the most abundant cell type. B) TREM2 and CD200 are jointly enriched in both secretome and lysate in microglia or neurons, respectively. In C) two examples, of proteins specifically enriched in the secretome, but not in the lysate are shown. ADIPOQ is specifically secreted from oligodendrocytes, but reveals highest expression in astrocytes. CACHD1 is specifically secreted from neurons, but high protein levels can be found in astrocytes, neurons and oligodendrocytes.

Because we observed that shedding contributes significantly to protein secretion, particularly in neurons, we considered the possibility that cell type-specific protein shedding may mechanistically also depend on cell type-specific expression of the contributing shedding protease. For example, the Alzheimer’s disease-linked protease *β*-site APP cleaving enzyme (BACE1)(Vassar, Bennett et al., 1999, Yan, Bienkowski et al., 1999), which has fundamental functions in the brain, is highly expressed in neurons, but not in astrocytes, microglia or oligodendrocytes (Voytyuk, Mueller et al., 2018). Consistent with our hypothesis, several known BACE1 substrates (APLP1, CACHD1, PCDH20 and SEZ6L) which are broadly expressed, were specifically shed from neurons (examples shown in Fig. 4A,B and Supplementary Fig.5). To investigate whether additional proteins in our secretome resource may be shed by BACE1 in a cell type-specific manner, primary neurons were treated with the established BACE1 inhibitor C3 (Stachel, Coburn et al., 2004)(Fig. 5A; Supplementary Table 6), followed by hiSPECS. This yielded 29 membrane proteins with reduced ectodomain levels in the secretome (Fig. 5B and Supplementary Fig.6A,B; Supplementary Table 7), which were scored as BACE1 substrate candidates. Besides known substrates, hiSPECS identified additional BACE1 substrate candidates (ADAM22, CD200, CXADR, and IL6ST) in neurons (Fig. 5B).

**Fig. 5:**
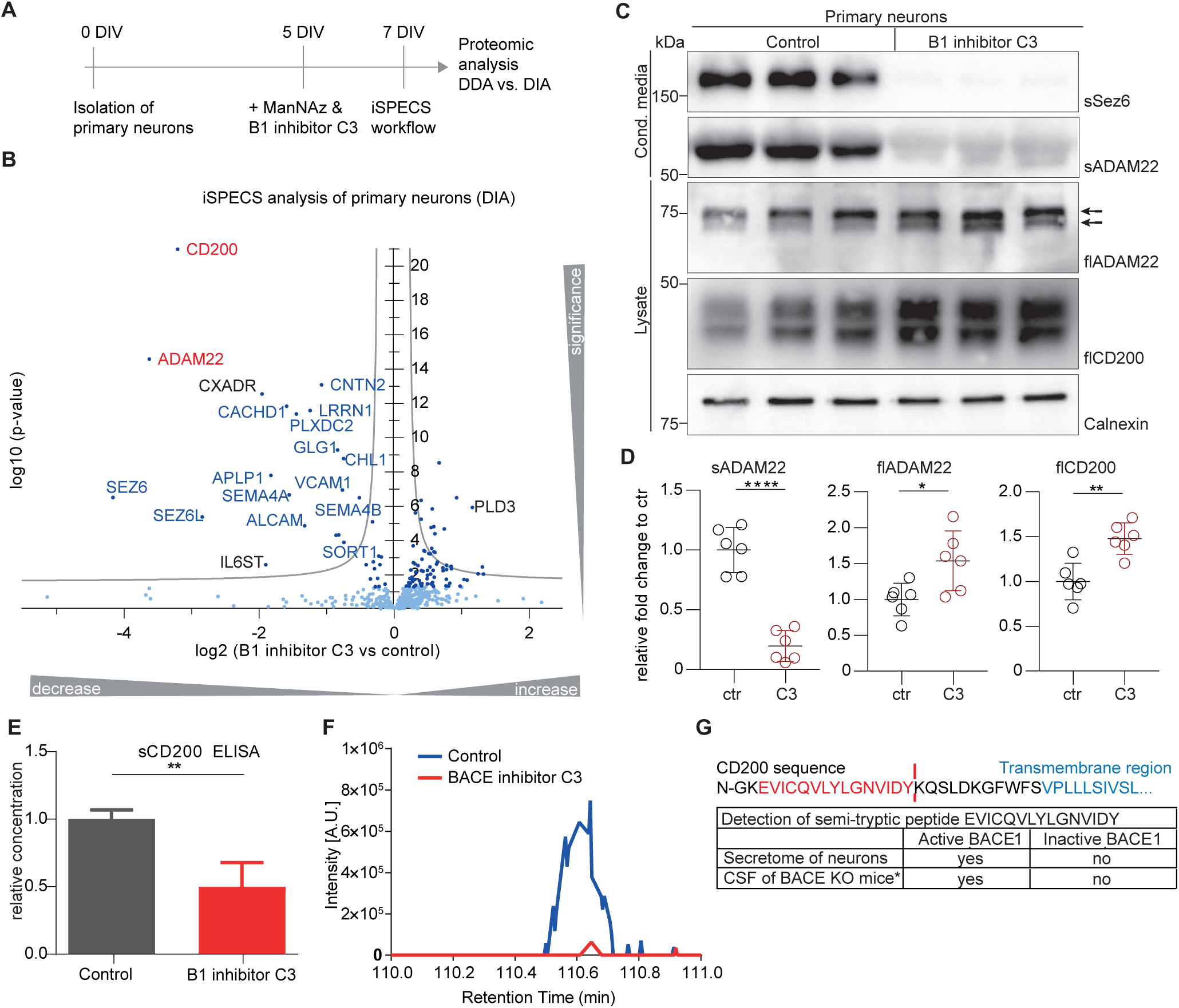
Identification and validation of substrate candidates of the protease BACE1. **A)** Experimental design of the ManNAz labeling step and BACE1 inhibitor C3 treatment of the primary neuronal culture for 48h at 5 to 7 days in vitro (DIV). **B)** Volcano plot showing changes in protein levels in the secretome of primary neurons upon BACE1 inhibitor C3 treatment using the hiSPECS DIA method. The negative log10 p-values of all proteins are plotted against their log2 fold changes (C3 vs control). The grey hyperbolic curves depict a permutation based false discovery rate estimation (p = 0.05; s0 = 0.1). Significantly regulated proteins (p<0.05) are indicated with a dark blue dot and known BACE1 substrates are indicated with blue letters. The two newly validated BACE1 substrates CD200 and ADAM22 are indicated in red. **C)** Independent validation of the novel BACE1 substrate candidates CD200 and ADAM22 by Western blotting in supernatants and lysates of primary neurons incubated with or without the BACE1 (B1) inhibitor C3 for 48h. Full-length (fl) ADAM22 and CD200 levels in the neuronal lysate and were mildly increased upon BACE1 inhibition, as expected due to reduced cleavage by BACE1. Calnexin served as a loading control. The soluble ectodomain of ADAM22 (sADAM22) was strongly reduced in the conditioned medium upon BACE1 inhibition. Ectodomain levels of the known BACE1 substrate SEZ6 (sSEZ6) were strongly reduced upon BACE1 inhibition and served as positive control. **D)** Quantification of the Western blots in C). Signals were normalized to calnexin levels and quantified relative to the control (ctr) condition. Statistical testing was performed with N=6 biological replicates, using the one sample t-test with the significance criteria of p < 0.05. According to this criterion, ADAM22 and CD200 were significantly increased in total lysates upon C3 treatment (flADAM22: p-value 0.0251, flCD200: p-value 0.011). Soluble ADAM22 was significantly reduced in the supernatant upon C3 treatment (p-value <0.0001). **E)** The reduction of the soluble ectodomain of CD200 (sCD200) was detected by ELISA, because the available antibodies were not sensitive enough for Western blots of the conditioned medium. **F)** Extracted ion chromatogram of the semi-tryptic peptide of CD200 in conditioned media of neurons comparing C3-treated to control condition. Levels of the semi-tryptic peptide were strongly reduced upon BACE1 inhibition. **G)** The potential cleavage site of CD200 was identified by a semi-tryptic peptide indicated in red which is from the juxtamembrane domain of CD200. The transmembrane domain is indicated in blue. The semi-tryptic peptide was found i) using the hiSPECS method in the neuronal secretome under control conditions but not upon BACE1 inhibition, and ii) in the CSF of wildtype mice but not upon knockout of BACE1 and its homolog BACE2 – *data are extracted from (Pigoni et al., 2016). Because BACE2 is hardly expressed in brain, both data sets demonstrate that generation of the semi-tryptic peptide requires BACE1 activity and thus, represents the likely BACE1 cleavage site in CD200.

ADAM22, which is a new BACE1 substrate candidate, and CD200, which was previously suggested as a BACE1 substrate candidate in a peripheral cell line (Stutzer, Selevsek et al., 2013), were further validated as neuronal BACE1 substrates by Western blots and ELISA assays (Fig. 5C-E). For CD200 we also detected a semi-tryptic peptide in the conditioned medium of neurons and in a previous proteomic data set (Pigoni, Wanngren et al., 2016) of murine cerebrospinal fluid (CSF) (Fig. 5F,G), but not when BACE1 was inhibited in the neurons or in the CSF of mice lacking BACE1 and its homolog BACE2 (Pigoni et al., 2016). As this semi-tryptic peptide derives from the juxtamembrane domain of CD200 where BACE1 typically cleaves its substrates, this peptide likely represents the cleavage site of BACE1 in CD200 (Fig. 5G). The validation of CD200 as a new *in vivo* BACE1 substrate demonstrates the power of hiSPECS to unravel the substrate repertoire of transmembrane proteases and reveals that cell type-specific expression of an ectodomain shedding protease is an important mechanism controlling the cell type-specific secretion/shedding of proteins.

Taken together, cell type-specific protein secretion and shedding is minimally dependent on cell type-specific protein expression of the secreted protein. Instead, cell type-specific expression of shedding regulators and other mechanisms to be discovered have an important role in determining cell type-specific protein secretion.

### Cell type-specific origin of CSF proteins

Cerebrospinal fluid (CSF) is the body fluid in direct contact with the brain and serves as an *in vivo* secretome. It is widely used for basic and preclinical research as well as clinical applications. However, a major limitation of CSF studies is that changes in CSF composition induced by disease or treatments often cannot be traced back to the cell type of origin, because most CSF proteins are expressed in multiple cell types (Sharma et al., 2015). To overcome this limitation, we applied the cell type-resolved brain secretome resource derived from the four most abundant brain cell types to determine the likely cellular origin of secreted glycoproteins in CSF (Supplementary Table 8).

In CSF from individual wild-type mice, 984 protein groups were identified with DIA in at least 3 out of 4 biological replicates (Fig. 6A), which represents higher coverage than in previous mouse CSF studies (Dislich, Wohlrab et al., 2015, Pigoni et al., 2016) and underlines the superiority of DIA over DDA in samples with low protein concentration. Proteins were grouped according to their log_10_ DIA LFQ intensities (as a rough estimate of protein abundance) into quartiles and analyzed according to their UniProt annotations for membrane, secreted, and cytoplasm (Fig. 5A). Soluble secreted proteins, such as APOE, were more abundant in the 1^st^ quartile, whereas the shed ectodomains, e.g. of CD200 and ADAM22, were the largest group of proteins in the 3^rd^ and 4^th^ quartile, indicating their lower abundance in CSF as compared to the soluble secreted proteins.

**Fig. 6:**
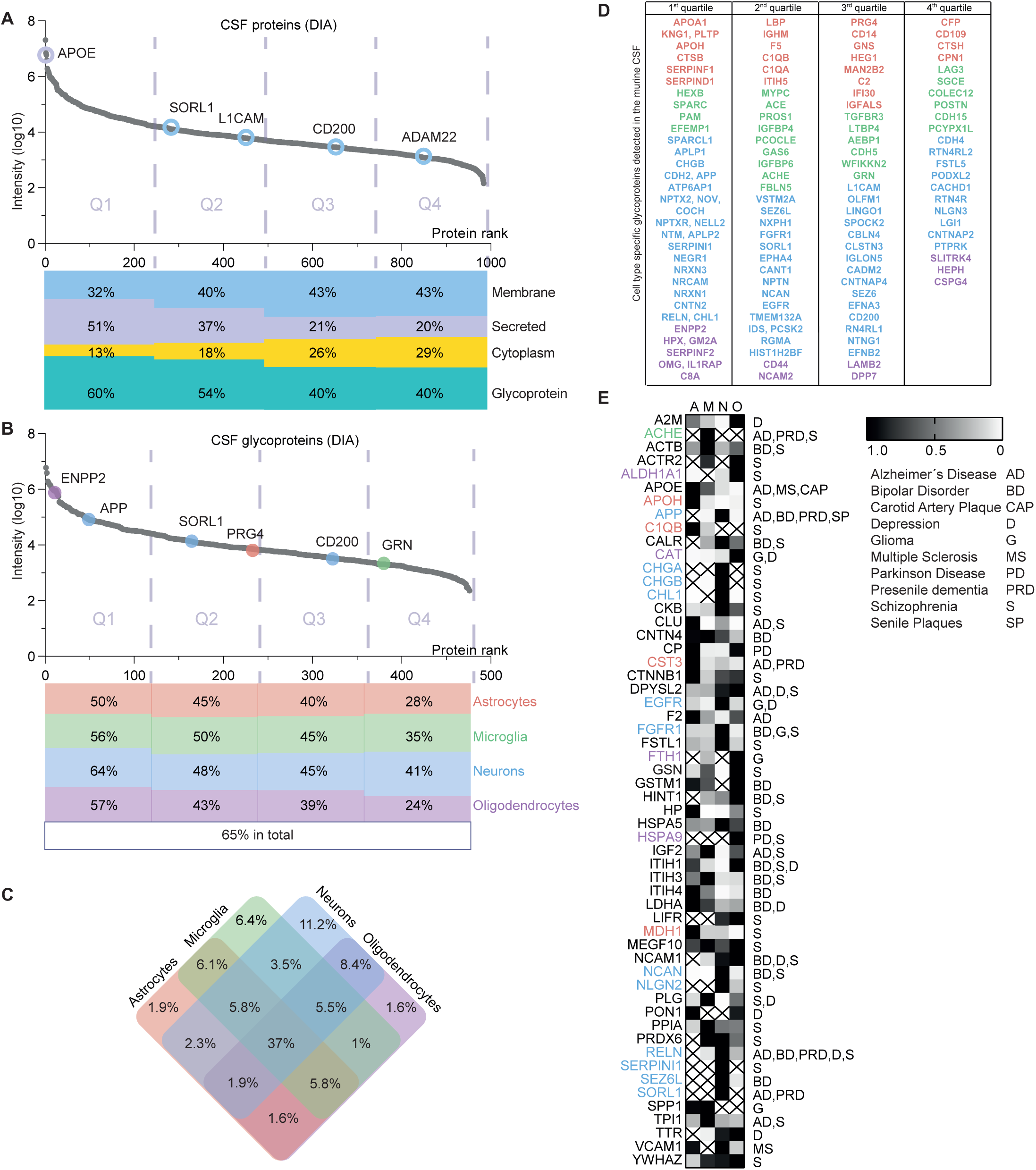
Mapping of murine CSF proteins to their probable cell type origin. **A)** Protein dynamic range plot of the log10 transformed LFQ intensities of the murine CSF proteins quantified in at least 3 of 4 biological replicates measured with DIA. The proteins are split into quartiles according to their intensities, with the 1st quartile representing the 25% most abundant proteins. The percentage of proteins annotated in UniProt with the following subcellular locations/keywords are visualized for: membrane, secreted, cytoplasm, and glycoprotein. **B)** Protein dynamic range plot of the log10 transformed LFQ intensities specifically of glycoproteins in the murine CSF quantified in at least 3 of 4 biological replicates measured with DIA. The proteins are split into quartiles according to their intensities. The percentage of proteins identified in the secretome of astrocytes, microglia, neurons or oligodendrocytes in at least 5 of 6 biological replicates is illustrated below. Selected proteins specifically secreted from one cell type are indicated with the color code of the corresponding cell type. **C)** Venn diagram indicating the distribution of CSF glycoproteins detected in the hiSPECS secretome resource. **D)** Cell type specifically secreted proteins (CTSP) in the hiSPECS secretome study (5-fold enriched in pairwise comparison to the other cell types or only detected in one cell type) are listed according to their presence in the CSF quartiles. **E)** List of proteins detected in murine CSF and the hiSPECS secretome resource which have human homologs that are linked to brain disease based on the DisGeNet database(Pinero et al., 2017) with an experimental index, e.i >= 0.9. Relative protein levels in the brain cell secretome is indicated (black high, white low abundance). Colored gene names indicate cell type-specific secretion.

Among the 476 CSF proteins annotated as glycoproteins (UniProt), 311 (65%) were also detected in the hiSPECS secretome resource and were mapped to the corresponding cell type (Fig. 6B). The most abundant glycoproteins of each cell type (top 25) revealed also high coverage in the CSF up to 76% of the top 25 neuronal proteins (Supplementary Fig. 7A). In general, proteins of neuronal origin represented the largest class of CSF glycoproteins and this was independent of their abundance (Fig. 6B,C). Given that astrocytes and oligodendrocytes outnumber neurons by far in the brain, this demonstrates that neurons disproportionately contribute to the CSF proteome. This prominent role of neurons is also reflected when focusing on the 420 cell type-specifically secreted proteins of our brain secretome resource. 123 of these proteins were also found in murine CSF (Fig. 6D). The majority in each quartile were proteins of neuronal origin. Similar to the proteins secreted from the four major brain cell types *in vitro*, the largest amount of the cell type-specifically secreted proteins detected in CSF are expressed in multiple cell types, but only secreted from one (Supplementary Table 4). This includes BACE1 substrates, such as SEZ6L and CACHD1, in agreement with BACE1 being expressed in neurons, but not in other brain cell types (Voytyuk et al., 2018). Thus, cell type-specific protein expression as well as cell type-specific protease expression and potentially additional mechanisms govern cell type-specific protein secretion, both *in vitro* (hiSPECS resource) and *in vivo* (CSF).

Several of the detected murine CSF proteins have human homologs linked to brain diseases (DisGeNET database (Pinero, Bravo et al., 2017)) and may serve as potential biomarkers. We mapped these proteins to their likely cell type of origin (Fig. 6E). Numerous proteins were specifically secreted from only one cell type, such as APP from neurons or granulin (GRN) from microglia, which have major roles in neurodegenerative diseases (Chitramuthu, Bennett et al., 2017, O’Brien & Wong, 2011). Thus, assigning the disease-related CSF/secretome proteins to one specific cell type offers an excellent opportunity to study the relevant cell type with regard to its contribution to disease pathogenesis. One example is the protein SORL1, which is genetically linked to Alzheimer’s disease (Yin, Yu et al., 2015). Although SORL1 is similarly expressed in all four major brain cell types, it was specifically released from neurons in the hiSPECS resource (Supplementary Table 4), indicating that pathology-linked changes in CSF SORL1 levels are likely to predominantly result from neurons.

Large-scale proteomic analyses of AD brain tissue and CSF (Bai, Wang et al., 2020, Johnson et al., 2020) continue to reveal more AD-linked proteins, such as increased CD44 in the CSF of AD patients (Johnson et al., 2020). Our resource demonstrates that CD44 is predominantly secreted from oligodendrocytes among the mouse brain cells, although having similar protein abundance in different brain cell types (Sharma et al., 2015)(Supplementary Fig.5). This data suggest to focus on oligodendrocytes for future studies determining how increased CD44 levels are mechanistically linked to AD pathophysiology. Taken together, the hiSPECS resource enables systematic assignment of CSF glycoproteins to the specific cell type of origin, which offers multiple opportunities to study CNS diseases.

### Cell type-resolved brain tissue-secretome

Next, we tested whether hiSPECS and the brain secretome resource can be used to determine the cell type-resolved secretome of brain tissue. We used organotypic cortico-hippocampal brain slices (Supplementary Table 9), an *ex-vivo* model of the brain (Daria, Colombo et al., 2017) that preserves the complex network of the diverse brain cell types. Despite the high amounts (25%) of serum proteins, 249 protein groups were identified in at least 5 of 6 biological replicates using hiSPECS DIA (Fig. 7A), demonstrating that hiSPECS is applicable for *ex vivo* brain tissue. Proteins were grouped according to their log_10_ abundance into quartiles (Fig. 7B). Similar to CSF, the majority of the more abundant proteins in the 1^st^ and 2^nd^ quartile were soluble proteins, whereas the shed transmembrane protein ectodomains were in the 3^rd^ and 4^th^ quartile, indicating their lower protein abundance compared to the soluble secreted proteins. 89% of the proteins detected in the slice culture secretome were also detected in the hiSPECS secretome library of the different brain cell types, which allows tracing them back to their cellular origin (Fig 7B). The cell-type resolved tissue-secretome revealed on average the highest contribution of microglia (74%), followed by astrocytes (72.8%), oligodendrocytes (68.5%) and neurons (66.5%). Interestingly, in quartile one, which resembles the most abundant proteins, oligodendrocytes are the most prominent with 85%. In addition, numerous cell type-specifically secreted proteins were identified (Fig. 7C).

**Fig. 7:**
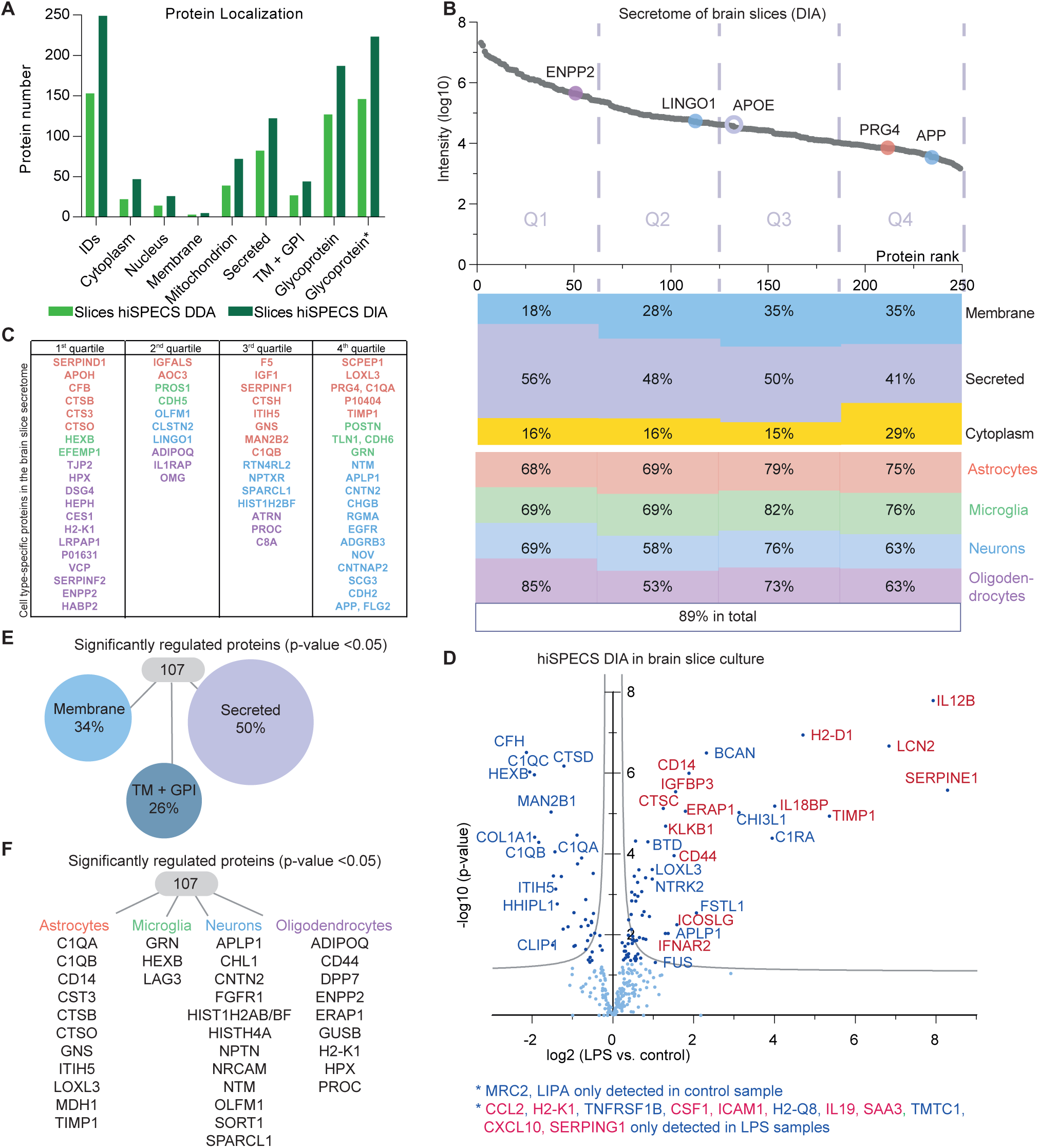
The secretome of brain slices. **A)** hiSPECS DDA and DIA analysis of brain slice cultures in the presence of 25% serum. The bar chart comparing the hiSPECS DDA and DIA method indicates the protein number and their localization according to UniProt identified in the secretome of brain slices. Proteins quantified in at least 5 of 6 biological replicates are considered. **B)** Protein dynamic range plot of the log10 LFQ intensities of secretome proteins of brain slices in descending order. According to their intensity, proteins are grouped into quartiles and the percentage of proteins with the following UniProt keywords is visualized: membrane, secreted, cytoplasm. The percentage of proteins identified in the secretome of astrocytes, microglia, neurons or oligodendrocytes in at least 5 of 6 biological replicates is illustrated. Selected proteins specifically secreted from one cell type are indicated with the color code of the corresponding cell type. **C)** Cell type-specifically secreted proteins according to the hiSPECS secretome resource (5-fold enriched in pairwise comparison to the other cell types or consistently detected only in one cell type; Supplementary Table 4) detected in the secretome of brain slices. **D)** Volcano plot showing changes in protein levels in the secretome of primary cultured brain slices upon 6 h LPS treatment in a 24 h collection window using the hiSPECS DIA method. The negative log10 p-values of all proteins are plotted against their log2 fold changes (LPS vs control). The grey hyperbolic curves depict a permutation based false discovery rate estimation (p = 0.05; s0 = 0.1). Significantly regulated proteins (p<0.05) are indicated with a dark blue dot. Proteins highlighted in red indicate proteins known to increase upon LPS treatment(Meissner et al., 2013) for details see Supplementary Table 9. **E-F)** Significantly regulated proteins upon LPS treatment (p<0.05) of brain slicese) split according to UniProt keywords membrane, single-pass transmembrane and GPI-anchored, and secreted proteins; F) indicating cell type specificity in the hiSPECS secretome resource.

As a final application of the brain secretome resource, we treated brain slices for 6 h with the strong inflammatory stimulus lipopolysaccharide (LPS), which serves as a model for acute neuroinflammation. Conditioned medium was analyzed using hiSPECS DIA. LPS strongly changed the brain slice secretome. Secretion of several proteins (marked in red) known to be LPS-responsive in macrophages(Meissner et al., 2013), such as H2-D1, CD14, or IL12B, were strongly upregulated (up to 250-fold) (Fig. 7D, Supplementary Table 9). Likewise, secretion of several proteins not known to be secreted in an LPS-dependent manner, such as BCAN, CHI3L1 and the complement receptor C1RA, were strongly upregulated. Among the 107 significantly regulated proteins (p<0.05) 50% and 26% are annotated as secreted proteins or single pass transmembrane proteins, respectively (Fig. 7E). Comparison of the brain slice secretome data to the secretome resource revealed that several proteins were secreted cell type-specifically, such as SORT1 and CHL1 by neurons and ENPP5 and ADIPOQ by oligodendrocytes (Fig. 7F). This demonstrates that not only immune cells responded with changes in their secretome to the inflammation cue, but instead indicates a systemic inflammatory response of multiple cell types. Moreover, this experiment suggests that systematic, proteome-wide secretome analysis of *ex vivo* brain slices is well suited to identify proteins and cell types contributing to neuroinflammation and potentially neurodegeneration.

## Discussion

Omics’ approaches have generated large collections of mRNA and protein abundance data across different cell types, including microglia and neurons, but we know very little about the molecules that are secreted from cells. This information is essential for our understanding of basic mechanisms of protein secretion, cell-cell communication within organs, particularly within the brain, and for identification of suitable biomarkers for brain processes in health and disease, such as APOE and TREM2 in AD(Huang, Zhou et al., 2019, Wolfe, Fitz et al., 2018).

Our development of the novel method hiSPECS miniaturizes mass spectrometry-based secretome analysis and enables secretome analysis of primary cell types, including the lower abundant ones in the brain. Importantly, hiSPECS allows culturing cells in the presence of serum or physiological cell culture supplements, whereas most previous secretome studies were restricted to serum- and even protein-free culture conditions, which is not feasible with many primary cell types. Besides its broad applicability to primary cells, we demonstrate that hiSPECS can also be applied to brain tissue *ex vivo*, which is widely used in neuroscience. The streamlined hiSPECS workflow also facilitates a broad applicability in laboratories without proteomic expertise. The strongly reduced mass spectrometer measurement time enables cost-effective, single-shot proteomic analysis of the samples.

With hiSPECS we established the cell type-resolved secretome resource of the four major cell types in the brain and its application to map the putative cell type-specific origin of CSF proteins (*in vivo* secretome) and proteins secreted from brain slices (*ex vivo* secretome). This approach provided fundamental new biological insights. First, ectodomain shedding is a major mechanism contributing to the protein composition of the secretome and quantitatively differs between brain cell types. Second, shed proteins *in vitro* (primary cells), *ex vivo* (brain slices) and *in vivo* (CSF) have a generally lower abundance in the secretome than secreted soluble proteins. This is consistent with the function of the shedding process as a regulatory mechanism, which releases bioactive membrane protein ectodomains on demand into the secretome, as seen with cytokines(Lichtenthaler et al., 2018). Thus, shedding provides an additional layer of control for the composition of the secretome that goes beyond constitutive protein secretion. Third, protein secretion is a highly cell type-specific process in the nervous system. This is surprising because we found that more than 73% of the cell type-specifically secreted proteins were expressed in more than one cell type, demonstrating that cells do not simply control secretion through cell type-specific expression of the secreted protein, but instead must have acquired additional mechanisms to control cell type-specific protein secretion, which are not yet well understood. Our resource provides insights into the underlying mechanisms. One example is the cell type-specific expression of a shedding protease, such as BACE1 in neurons. Ectodomain shedding happens for more than 1,000 membrane proteins (Lichtenthaler et al., 2018), but in most cases, the contributing protease is not known. Thus, it is likely that shedding proteases other than BACE1 are also expressed in a cell type-specific manner, and thus contribute to cell type-specific protein shedding in the brain.

While additional mechanisms underlying cell type-specific protein secretion remain to be elucidated, protein transport through the secretory pathway, which is a prerequisite for protein secretion or shedding, is a potential mechanism of regulation. In fact, some soluble proteins require CAB45 for their exit from the trans-Golgi network, whereas other proteins do not(Blank & von Blume, 2017). Likewise, some transmembrane proteins require specific transport helpers for trafficking through the secretory pathway such as IRHOM1/2 for ADAM17 and Cornichon for transforming growth factor(Dancourt & Barlowe, 2010, Lichtenthaler, 2012). Thus, it appears possible that transport-selective proteins may be preferentially expressed in some brain cell types and consequently allow for a cell type-specific secretion or shedding of their cargo proteins. hiSPECS is an excellent method to address these fascinating questions regarding the complex mechanisms controlling protein secretion in the nervous system.

The secretome of the four major brain cell types and the *ex vivo* tissue identified here represents a snapshot of the total brain secretome and more secreted proteins are known or likely to exist. These include non-glycosylated secreted proteins which are not captured with hiSPECS as well as proteins secreted in a time-dependent manner such as during development or aging. For example, it is known that microglia can partially change their expression profile when taken into the culture, which may consequently affect the secretome (Gosselin, Skola et al., 2017). Additionally, the secretome may change upon cell stimulation, such as during neuronal activity or inflammation or when different cell types are cocultured or taken into three-dimensional culture systems (Stiess, Wegehingel et al., 2015). Additional proteins may be selectively secreted from lower abundant brain cell types, such as pericytes.

Taken together, hiSPECS and the cell type-resolved mouse brain secretome resource are important new tools for many areas in neuroscience, from mechanisms of protein secretion and signal transduction between brain cells *in vitro*, *ex vivo* (brain slices) and *in vivo* (CSF) to functional analysis of nervous system proteins (identification of protease substrates) and cell type-specific biomarker determination in CSF (e.g. CD44) with high relevance to psychiatric, neurological and neurodegenerative diseases.

## Acknowledgement

We thank Felix Meissner and Jürgen Cox for helpful comments on this manuscript, and Anna Daria for support with the *ex vivo* model. This work was supported by the Deutsche Forschungsgemeinschaft (German Research Foundation) within the framework of the Munich Cluster for Systems Neurology (EXC 2145 SyNergy, project ID 390857198) and the BMBF through project CLINSPECT-M and JPND PMG-AD. JT was supported by a Boehringer Ingelheim Fonds (BIF) PhD fellowship. Funds have been provided to ST by the Alzheimer Forschung Initiative e.V. (project number 18014)

## Author Contributions

JT, SAM, SFL designed the experiments and prepared the manuscript. JT, SAM performed the proteomic analysis of all samples. JT, SAM analyzed the data with bioinformatics input from ESK, JZ, DF and GG. LSM, ST provided primary microglia and cortico-hippocampal brain slices. MS and MS provided primary oligodendrocyte culture and MS also gave conceptual advice.

## Materials and Methods

### Data Availability

The proteomic resource is available for the public on the Proteome Xchange Consortium via the PRIDE Archive (Project accession: PXD018171).

### Mice

All murine samples were isolated from C57BL/6J mice from The Jackson Laboratory according to the European Communities Council Directive. Mice were housed and breed in the pathogen free animal facility of the DZNE Munich.

### Primary cell culture and brain slices

All primary cultures were maintained under standard cell culture conditions at 37 °C with 5% CO_2_. The samples were collected from at least two independent culture preparations, with 3 dishes of primary cells from each culture preparation. It is not possible to determine the sex of the cells, because the cells were isolated from embryos or young pups. The conditioned media were stored at −20 °C until further processing.

Primary neurons were isolated at E16.5 as described before(Kuhn et al., 2012). Meninges-free cerebral cortices or hippocampi were dissociated and digested in DMEM with 200 u Papain for 30 min (Sigma Aldrich) and plated into poly-D-lysine coated 6-wells in plaiting media (DMEM + 10% FBS + 1% penicillin/streptomycin). After 4h media was changed to neuronal cultivation media (B27 + Neurobasal + 0.5 mM glutamine + 1% P/S).

Primary astrocytes were isolated at E16.5 and dissociated in the same way as the primary neurons, however, they were plated on uncoated dishes. Cultures were grown until reaching confluence in DMEM + 10% FBS, cells were detached using trypsin and re-seeded on a new plate (50% confluence). This procedure was repeated three times before seeding the cells for the final experiment (1×10^6^ cells into a well of a 6-well plate).

Primary oligodendrocyte progenitor cell (OPC) cultures were prepared by magnetic-activated cell sorting (MACS). 60 mm cell culture dishes were coated with 0.01% poly-L-lysine for 1 h at 37 °C, washed twice, and incubated with MACS Neuro Medium (Miltenyi Biotec, #130-093-570) overnight. OPCs were isolated from the brains of postnatal day 6 C57BL/6J mouse pups. Cell suspension was obtained by automated dissociation using the Neural Tissue Dissociation Kit (P) (Miltenyi Biotec, Cat#130-092-628) and the gentleMACS™ Dissociator (Miltenyi Biotec, Cat#130-093-235) following the datasheet of the kit with some modifications. DMEM/pyruvate medium was used instead of HBSS during tissue dissociation. All media were warmed up to room temperature. The optional centrifugation steps were included in the dissociation. The myelin removal step was omitted. Prior to labeling with anti-AN2 MicroBeads (Miltenyi Biotec, Cat#130-097-170), the cell suspension was incubated with the OPC MACS cultivation medium (MACS Neuro Medium containing MACS NeuroBrew-21 (Miltenyi Biotec, #130-093-566), GlutaMAX (Thermo Fisher Scientific, #35050087) and penicillin/streptomycin) for 3h at 37 °C for surface antigen re-expression. DMEM containing 1% horse serum and penicillin/streptomycin (DMEM/HS medium) was used as the buffer for magnetic labeling and separation. After magnetic separation, the OPC MACS cultivation medium was applied to flush out AN2^+^ cells. 1×10^6^ cells were plated in 4 mL of OPC MACS cultivation medium per 60mm dish. For OPC/oligodendrocyte culture, ManNAz was added directly to the medium after one day *in vitro* and incubated for 48h

Primary microglia were isolated from postnatal day 5 mouse brains using the MACS technology as previously described(Daria et al., 2017) Briefly, olfactory bulb, brain stem and cerebellum were removed and the remaining cerebrum was freed from meninges and dissociated by enzymatic digestion using the Neural Kit P (Miltenyi Biotec; Cat#130-092-628). Subsequently, tissue was mechanically dissociated by using three fire-polished glass Pasteur pipettes of decreasing diameter. Microglia were magnetically labelled using CD11b MicroBeads (Miltenyi Biotec, Cat# 130-093-634) and cell suspension was loaded onto a MACS LS Column (Miltenyi Biotec) and subjected to magnetic separation. 1.5-2x 10^6^ microglia were then cultured in DMEM/F12 media (Invitrogen) supplemented with 10% heat inactivated FBS (Sigma) and 1% Penicillin-Streptomycin (Invitrogen) in a 60 mm dish for four days before the 48h treatment with ManNAz. Conditioned media of 1×10^6^ cells was used for the hiSPECS experiment.

Organotypic brain slice cultures from young (postnatal day 6-7) mice were prepared as described previously(Daria et al., 2017). Briefly, after brain isolation, olfactory bulb, midbrain, brain stem and cerebellum were removed and the two remaining cortical hemispheres cut at 350 µm with a tissue chopper (Mcllwain, model TC752, Mickle Laboratory Engineering Company). Intact sagittal cortico-hippocampal slices were selected and incubated for 30 min at 4°C in a pre-cooled dissection media (50% HEPES-buffered MEM, 1% penicillin–streptomycin, 10 mM Tris, pH 7.2). Four slices were then plated onto a 0.4-um porous polytetrafluoroethylene (PTFE) membrane insert (PICMORG50, Millipore) placed in a 35 mm dish with a slice culture media containing 50% HEPES-buffered MEM, 25% HBSS, 1 mM L-glutamine (Gibco) and 25% heat-inactivated horse serum (Merck-Sigma). Media was exchanged one day after preparation and subsequently every three days. Brain slices were cultured for 14 days before the 48h treatment with ManNAz.

### hiSPECS

After washing the primary cells with 1x PBS, cell-type specific growth media containing serum supplements with 50 µM of ManNAz (Thermo Fisher Scientific, Cat#C33366) was added for 48h. Afterwards, conditioned media was collected and filtered through Spin-X 0.45 µM cellulose acetate centrifuge tube filter (#8163, Costar) and stored at −20°C in protein Lobind tubes (Eppendorf) until further usage. Glycoprotein enrichment was performed using 60 µL Concanavalin A (ConA) bead slurry per sample (Cat#C7555, Sigma Aldrich). ConA beads were washed twice with 1 mL of binding buffer (5 mM MgCl_2_, 5 mM MnCl_2_, 5 mM CaCl_2_, 0.5 M NaCl, in 20 mM TrisHCL pH 7.5) before use. The conditioned medium was incubated with the ConA beads for 2h in an overhead rotator at room temperature. The ConA beads were pelleted by centrifugation (2000 g, 1 min) and the supernatant containing unbound proteins was discarded. The beads were washed three times with 1 mL binding buffer before adding 500 µL of elution buffer (500 mM Methyl-alpha-D-mannopyranoside, 10 mM EDTA in 20 mM TrisHCl pH 7.5) and rotating over-head for 30 min at RT. The eluate was filtered through pierce spin columns (Thermo, #69725) to remove remaining ConA beads and then the filtrate was transferred to a 1.5 mL protein Lobind tube. The elution step was repeated with another 500 µL elution buffer and combined with the first eluate. 50 µL of magnetic DBCO beads (Jena bioscience, Cat#CLK-1037-1) were washed twice with mass spec grade water and added to the eluate. Sodium deoxycholate (SDC) was added to a final concentration of 0.1% (w/v) to prevent clumping/sticking of the beads (except otherwise noted).The click reaction was performed overnight at 4°C on an Eppendorf Thermomixer R shaker at 1200 rpm to covalently couple metabolically labeled glycoproteins to the magnetic beads. The next day, beads were washed three times with 1 mL 1% SDS buffer (100mM TrisHCl pH 8.5, 1% SDS, 250 mM NaCl), three times with 1 mL 8 M UREA buffer (8 M Urea in 100mM TrisHCl pH 8.5) and three times with 1 mL 20% (v/v) acetonitrile. Beads were retained with a magnetic rack (Dynamag-2, Thermo Scientifc). After each step, samples were resuspended briefly by shaking at 1200 rpm at room temperature. Beads were transferred to a new 1.5 mL low binding tube using 2 x 500 µL mass spec grade water. Beads were retained in a magnetic rack and the supernatant was removed. Protein disulfide bonds were reduced in 50 µL of 10 mM dithiothreitol (DTT) in 100 mM ammonium bicarbonate (ABC) for 30 min at 1200 rpm at 37°C. Afterwards, the supernatant was discarded. Alkylation of cysteines was performed using 50 µL of 55 mM iodoacetamide (IAA) in 100mM ABC for 30 min at 1200 rpm and 20°C in the dark. The supernatant was discarded and beads were washed twice with 100 µL of 100 mM ABC. The protein digestion was performed by adding 0.2 µg LysC (Promega) in 50 µL of 100 mM ABC for 3h at 1200 rpm at 37°C followed by overnight trypsin digestion using 0.2 µg of trypsin (Promega) per sample in 100 mM ABC without 0.1% SDC. The supernatant containing the tryptic peptides was transferred to a 0.5 mL protein Lobind tube. Beads were washed twice with 100 mM ABC without 0.1% SDC and added to the same tube. Each sample was acidified with 50 µL of 8% FA and incubated for 20 min at 4°C. Precipitated SDC was removed by centrifugation at 18.000 g for 20 min at 4 °C. Peptides were cleaned up using C18 Stage tips as previously described (Rappsilber, Ishihama et al., 2003). Dried peptides were resuspended in 18 µL 0.1% formic acid (FA) and 2 µL of 1:10 diluted iRT peptides (Biognosys, Ki-3002-1) were spiked into the samples.

### Mass spectrometry

The LC-MS/MS analyses were performed on an EASY-nLC 1200 UHPLC system (Thermo Fisher Scientific) which was online coupled with a NanoFlex ion source equipped with a column oven (Sonation) to a Q Exactive™ HF Hybrid Quadrupol-Orbitrap™ mass spectrometer (Thermo Fisher Scientific). 8 µL per sample were injected. Peptides were separated on a 30 cm self-made C18 column (75 µm ID) packed with ReproSil-Pur 120 C18-AQ resin (1.9 µm, Dr. Maisch GmbH). For peptide separation, a binary gradient of water and 80% acetonitrile (B) was applied for 120 min at a flow rate of 250 nL/min and a column temperature of 50 °C: 3% B 0 min; 6% B 2 min; 30% B 92 min; 44% B 112 min; 75% B 121 min.

Data dependent acquisition (DDA) was used with a full scan at 120,000 resolution and a scan range of 300 to 1400 m/z, automatic gain control (AGC) of 3×10^6^ ions and a maximum injection time (IT) of 50 ms. The top 15 most intense peptide ions were chosen for collision-induced dissociation (CID) fragmentation. An isolation window of 1.6 m/z, a maximum IT of 100 ms, AGC of 1×10^5^ were applied and scans were performed with a resolution of 15,000. A dynamic exclusion of 120 s was used.

Data independent acquisition (DIA) was performed using a MS1 full scan followed by 20 sequential DIA windows with variable width for peptide fragment ion spectra with an overlap of 1 m/z covering a scan range of 300 to 1400 m/z. Full scans were acquired with 120,000 resolution, AGC of 5×10^6^ and a maximum IT time of 120 ms. Afterwards, 20 DIA windows were scanned with a resolution of 30,000 and an AGC of 3×10^6^. The maximum IT for fragment ion spectra was set to auto to achieve optimal cycle times. The m/z windows were chosen based on the peptide density map of the DDA run of a representative hiSPECS sample and optimized in a way that allowed the detection of 8 points per peak. The following window widths were chosen according to the peptide density map (Fig. 1B):

|  |  |  |  |  |  |  |  |  |  |  |  |  |  |  |  |  |  |  |  |  |
| --- | --- | --- | --- | --- | --- | --- | --- | --- | --- | --- | --- | --- | --- | --- | --- | --- | --- | --- | --- | --- |
| Window # | 1 | 2 | 3 | 4 | 5 | 6 | 7 | 8 | 9 | 10 | 11 | 12 | 13 | 14 | 15 | 16 | 17 | 18 | 19 | 20 |
| Window Width (m/z) | 85 | 40 | 30 | 28 | 26 | 25 | 24 | 24 | 24 | 24 | 25 | 27 | 28 | 29 | 34 | 38 | 46 | 61 | 151 | 352 |

The following libraries were used for the different DIA experiments of this manuscript using the search engine platform MaxQuant and Spectronaut Pulsar X version 12.0.20491.14:

| Library | Unique modified peptides | Unique peptide precursors | Protein groups | Used in: |
| --- | --- | --- | --- | --- |
| hiSPECS neuron | 4,585 | 5,957 | 646 | Supplementary Table 1+7 |
| hiSPECS brain cell types | 12,695 | 15,886 | 1,540 | Supplementary Table 2 |
| CSF mouse | 20,100 | 24,985 | 2,550 | Supplementary Table 8 |
| hiSPECS slices | 2,675 | 3,375 | 431 | Supplementary Table 9 |

### BACE1 inhibitor treatment

The BACE inhibitor C3 (β-secretase inhibitor IV, Calbiochem, Cat#565788, Sigma Aldrich) or DMSO vehicle control were added in parallel to the ManNAz, at a final concentration of 2 μM, to the neurons for 48h (Kuhn et al., 2012).

### LPS treatment of brain slices

Organotypic brain slices were cultured for 2 weeks after the isolation process, followed by a 48 h ManNAz labelling step. Next, the brain slices were treated for 6 h with LPS (500 ng/mL) followed by a 24 h collection window of the conditioned media. Serum supplements were not added during the LPS treatment and the collection period to maximize the inflammatory response. However, additional ManNAz was added during all steps.

### Antibodies

The following antibodies were used for Western Blotting: Mouse monoclonal anti-ADAM22 (UC Davis/NIH NeuroMab Facility Cat#75-093), Rat monoclonal anti-SEZ6 (Pigoni et al., 2016), Goat polyclonal anti-CD200 (R and D Systems Cat#AF2724,), Rabbit polyclonal anti-calnexin (Enzo Life Sciences Cat#ADI-SPA-860).

### ELISA

CD200 ectodomain levels in the supernatant of primary neurons were measured and quantified using the following ELISA kit according to the supplier’s manual: Mouse CD200 ELISA Kit (LSBio Cat#LS-F2868). A volume of 250 µL from the total of 1 mL undiluted media of 1.5 million primary cortical neurons cultured for 48h was used per technical replicate. The standard curve provided with the kit was used and neuronal media which was not cultured with cells, was used as a blank value.

### Western blot analysis

Cells were lysed in STET buffer (50 mM Tris, pH 7.5, 150 mM NaCl, 2 mM EDTA, 1% Triton X-100), incubated for 20 min on ice with intermediate vortexing. Cell debris as well as undissolved material was removed by centrifugation at 20,000 × g for 10 min at 4°C. The protein concentration was determined using the BC assay kit of lnterchim (UP40840A) according to the manufacturer’s instructions. Samples were boiled for 10 min at 95°C in Laemmli buffer and separated on self-cast 8%, 10% or 12% SDS-polyacrylamide gels. Afterwards, proteins were transferred onto nitrocellulose membranes using a BioRad Wet/Tank Blotting system. The membranes were blocked for 20 min in 5% milk in 1xPBS with 1% Tween, incubated overnight at 4°C with the primary antibody, 1h at room temperature with the secondary antibody and developed using an ECL prime solution (GE Healthcare, RPN2232V1).

### Cerebrospinal fluid (CSF) sample preparation

In solution digestion of 5 µL CSF samples was performed as previously described (Pigoni et al., 2016). Dried peptides were dissolved in 18 µL 0.1% FA and 2 µL 1:10 diluted iRT peptides.

### Quantification and statistical analysis

In general, statistical details can be found in the Figure legends including statistical tests and n number used.

### Raw data analysis of mass spectrometry measurements

DDA raw data were analyzed with MaxQuant (version 1.5.5.1 or 1.6.1.0) using the murine UniProt reference database (canonical, downloaded on: 17.01.2018 which consisted of 16,954 proteins) and the Biognosys iRT peptide database for label free quantification (LFQ) and indexed retention time spectral library generation. Default settings were chosen, however, the minimal peptide length was set to six. Two missed cleavages were allowed, carbamidomethylation was defined as a fixed modification, and N-termini acetylation as well as oxidation of methionines were set as variable modifications. The false discovery rate (FDR) was set to less than 1% for protein and peptide identifications. The results of the MaxQuant analysis were used to generate DIA spectral libraries of proteins in Spectronaut Pulsar X (Biognosys). Data generated with DIA were analyzed using Spectronaut Pulsar X (Biognosys) using the self-generated spectral libraries applying default settings: quantification on the MS2 level of the top N (1 to 3) peptide spectra and a FDR of 1%.

### Bioinformatics analysis: Data Pre-processing and Normalization

For the hiSPECS brain secretome resource, the raw dataset was filtered to retain only the proteins that were consistently quantified in at least 5 of the 6 biological replicates (5/6 or 6/6) in any of the four cell types. This yielded a total of 995 out of 1083 quantified proteins. The data were further processed using Perseus (Version 1.6.6.0). The LFQ values were log2 transformed and the Pearson correlation coefficients between all samples were determined. An imputation procedure was employed by which missing values are replaced by random values of a left-shifted Gaussian distribution (shift of 1.8 units of the standard deviation and a width of 0.3).

### PCA and UMAP

Principal component analysis and uniform manifold approximation and projection (UMAP) (Diaz-Papkovich, Anderson-Trocme et al., 2019) were performed to visualize the relationships between the cell types and between the biological replicates. UMAP is a fast non-linear dimensionality reduction technique that yields meaningful organization and projection of data points. UMAP has the advantage of assembling similar individuals (or data points) while preserving long-range topological connections to individuals with distant relations.

### Differential abundance analysis

In order to determine differentially abundant proteins using pairwise comparisons of all four brain cell types we employed protein-wise linear models combined with empirical Bayes statistics (implemented in the R package Limma (Ritchie, Phipson et al., 2015), similarly to (Kammers, Cole et al., 2015)). A protein was considered as differentially abundant (DA) in the different brain cell types if the Bonferroni-corrected p-value was < 0.05 and the Log2 fold change ≥ 2. The Log2 fold change of 2 was set to reduce the number of false-positives due to data imputation.

### Pathway enrichment score

Proteins were considered to be mainly secreted from one cell type if proteins were identified in at least 5 out of 6 biological replicates and fulfilled one of following criteria: 1) proteins were only detected in two or less biological replicates in another cell type or 2) proteins were at least 5-fold enriched in a pairwise comparison to all three other cell types.

Functional annotation clustering of proteins specifically secreted from each cell type was performed with DAVID 6.8 (Huang da, Sherman et al., 2009a, Huang da, Sherman et al., 2009b) using all 995 robustly quantified proteins in the hiSPECS secretome resource as the background. The top three gene ontology terms for biological process (GOTERM_BP_FAT) were picked for visualization based on the enrichment score using medium classification stringency. The same settings were chosen to identify functional annotation clusters of all proteins in the hiSPECS library identified compared to the whole mouse proteome for the gene ontology term cellular compartment (GOTERM_CC_FAT).

### Comparison of secretome proteins to their relative abundances in the corresponding cell lysates using a published database

We compared our cell type-specific secretome proteins to the corresponding cell type-specifically expressed proteins identified in cell lysates of the same four cultured primary brain cell types in a previous proteomic study(Sharma et al., 2015). The data from their study containing the LFQ values of the individual biological replicates of the brain cell lysate were downloaded (*Sharma et al.*: Supplementary Table 1) and processed using Perseus. Proteins detected in at least one cell type with two biological replicates were considered and missing values were imputed as described before (replacement of missing values from left-shifted normal distribution, 1.8 units of the standard deviation and a width of 0.3). Proteins with 2.5-fold higher protein levels in one cell type as compared to all other three cell types were considered specific. On the basis of this analysis, we defined two categories: 1) proteins that were specifically enriched in one cell type both in cell lysates and the corresponding cell secretome and 2) cell type-specifically secreted proteins that were not enriched in the corresponding lysate. The second category indicates a possible secretome-specific mechanism, such as the selective secretion/shedding by a protease in the given cell type.

### Interactions with cell lysate proteins detected by Sharma et. al

To find the relationships between the proteins in the secretome and those from the cell lysate, we searched for interacting partners of the proteins specifically enriched in the secretome of a specific cell type and the proteins from the cell lysate as determined by (Sharma et al., 2015). We downloaded the mouse protein-protein interaction network (PPIN) from BioGRID (Chatr-Aryamontri et al., 2015) and additional binary interactions data from UniProt (UniProt Consortium, 2018).

### Disease association

To determine if the proteins significantly differentiated in one cell type with respect to the other 3 cell types are linked to neurodegenerative diseases, we searched for curated gene disease associations (GDA) from DisGeNET (Pinero et al., 2017). Our search list contained 31 known diseases of the nervous system. We set the evidence index (EI) to 0.95: an EI of 1 indicates that all the available scientific literature supports the specific GDA.

## Supplementary Information titles and legends

### Overview Supplementary Tables

Supplementary Table 1: Comparison of the SPECS and hiSPECS method using DIA and DDA. Related to Fig. 1.

Supplementary Table 2: hiSPECS secretome resources of astrocytes, microglia, neurons and oligodendrocytes. Related to Fig. 2.

Supplementary Table 3: Meta-analysis of brain cell lysate data and its comparison to the secretome resource. Related to Fig. 2,4.

Supplementary Table 4: Cell type-specifically secreted proteins of the hiSPECS secretome resource.

Related to Fig. 2-7.

Supplementary Table 5: Interaction analysis of cell type-specific secretome and transmembrane lysate proteins. Related to Fig. 3.

Supplementary Table 6: hiSPECS analysis of BACE1 inhibitor treated neurons vs. control.

Related to Fig. 5.

Supplementary Table 7: List of BACE1 substrate candidates.

Related to Fig. 5.

Supplementary Table 8: Murine CSF analysis and comparison to the cell type-resolved secretome resource. Related to Fig. 6.

Supplementary Table 9: hiSPECS analysis of brain slices.

Related to Fig. 7

## Supplemental Figure legends

**Supplementary Fig. 1:**
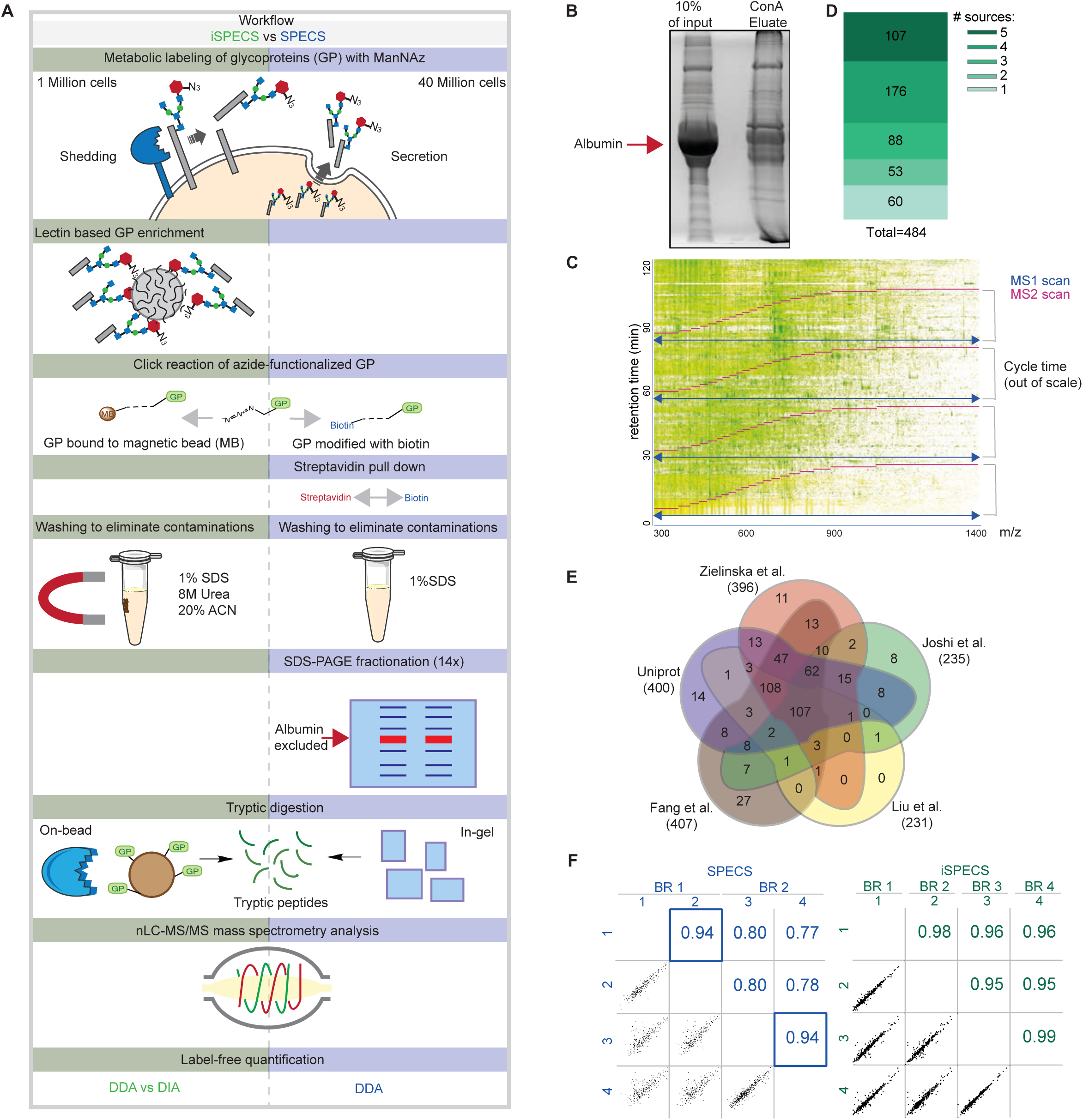
Benchmarking hiSPECS against SPECS. **A)** Comparison of the novel iSPECS (green) and the previous SPECS (blue) protocol (Kuhn et al., 2012). hiSPECS uses lectin-based glycoprotein enrichment followed by covalent binding to magnetic beads which improves sample processing. On-bead tryptic digestion is followed by mass spectrometry analysis and label-free quantification (LFQ) using data independent or data dependent acquisition (DIA vs DDA). In contrast, SPECS depends on biotinylation of the azide-functionalized glycoproteins, followed by streptavidin pull down, SDS-gel based fractionation into 14 gel slices and DDA analysis only. **B)** Coomassie stained gel showing the prominent reduction of albumin as a result of glycoprotein enrichment with ConA. Left lane: secretome (conditioned medium) of primary neurons before glycoprotein enrichment. 10% of the volume was loaded that was used for the ConA enrichment. Right lane: eluate after glycoprotein enrichment using ConA beads. The albumin band is highlighted with a red arrow. **C)** A representative peptide density plot of a neuronal secretome using the hiSPECS method and analyzed in DDA mode. The m/z ratio is plotted against the retention time. One MS1 scan (purple) is followed by 20 MS2 scans covering a range of 300 to 1400 m/z with an overlap of 1 m/z between adjoining m/z windows. The m/z windows were adjusted to achieve equal numbers of peptides. **D)** Identified glycoproteins in the secretome of primary neurons using the DIA hiSPECS protocol according to UniProt or (Fang et al., 2016, Joshi et al., 2018, Liu et al., 2017, Zielinska et al., 2010). **E)** Venn diagram illustrating in which sources the proteins of D) were found to contain a glycosylation site. As expected for the glyco-secretome, more than 85% of the quantified secretome proteins were annotated as glycoproteins. **F)** Representative Pearson correlations of log2 transformed protein LFQ intensities of four biological replicates (BR) either processed with the SPECS or hiSPECS method. In the previous SPECS studies sample pairs were separated on the same gel to achieve high reproducibility, whereas the correlation of biological replicates run on different gels was rather low. Thus, protein LFQ ratios of the individual replicates were used for statistical evaluation. The blue squares indicate samples run on the same gel during sample preparation (SPECS data obtained from (Kuhn et al., 2012)).

**Supplementary Fig. 2:**
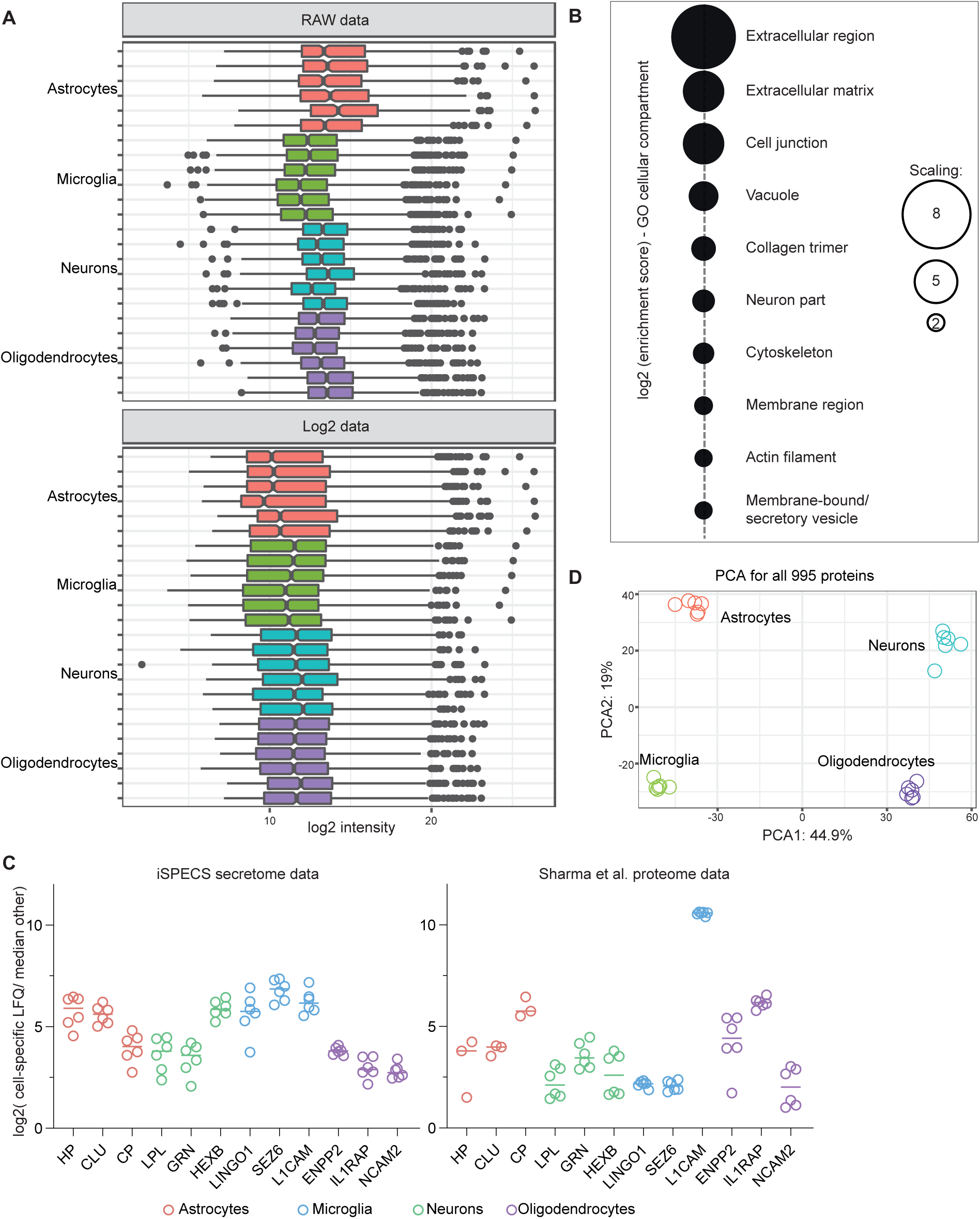
Quality control of cell type-resolved mouse brain secretome resource. **A)** Data transformation of the hiSPECS DIA secretome analysis of the brain cell types. Log2 Transformation of the label free quantification values (LFQ). **B)** Functional annotation clustering with DAVID 6.8 (Huang da et al., 2009a, Huang da et al., 2009b) for gene ontology term cellular component (FAT) of the 995 hiSPECS proteins identified using the mouse proteome as background. The dot sizes indicate the log 2 enrichment score. **C)** Fold change of cell type specific proteins in the brain cells. Log2 ratio of the average abundance in the specific cell type to the median abundance in the other cell types in the hiSPECS secretome or lysate analysis (Sharma et al., 2015) is shown. Known cell type-specific marker proteins are highlighted for each cell type which reveal a strong enrichment in the lysate and secretome of the primary brain cells verifying the quality and comparability of the primary cultures. For example, the ectodomain of the membrane protein NCAM2 (with an y-axis value of 3 in the log2 scale) is secreted about 8-fold more from oligodendrocytes compared to the median of the other three cell types. **D)** Principal component analysis (PCA). The secretomes of the four cell types segregated based on the two major components of all 995 proteins identified in at least 5 biological replicates in one cell type, which accounted for 44.9% and 19% of the variability, respectively.

**Supplementary Fig. 3:**
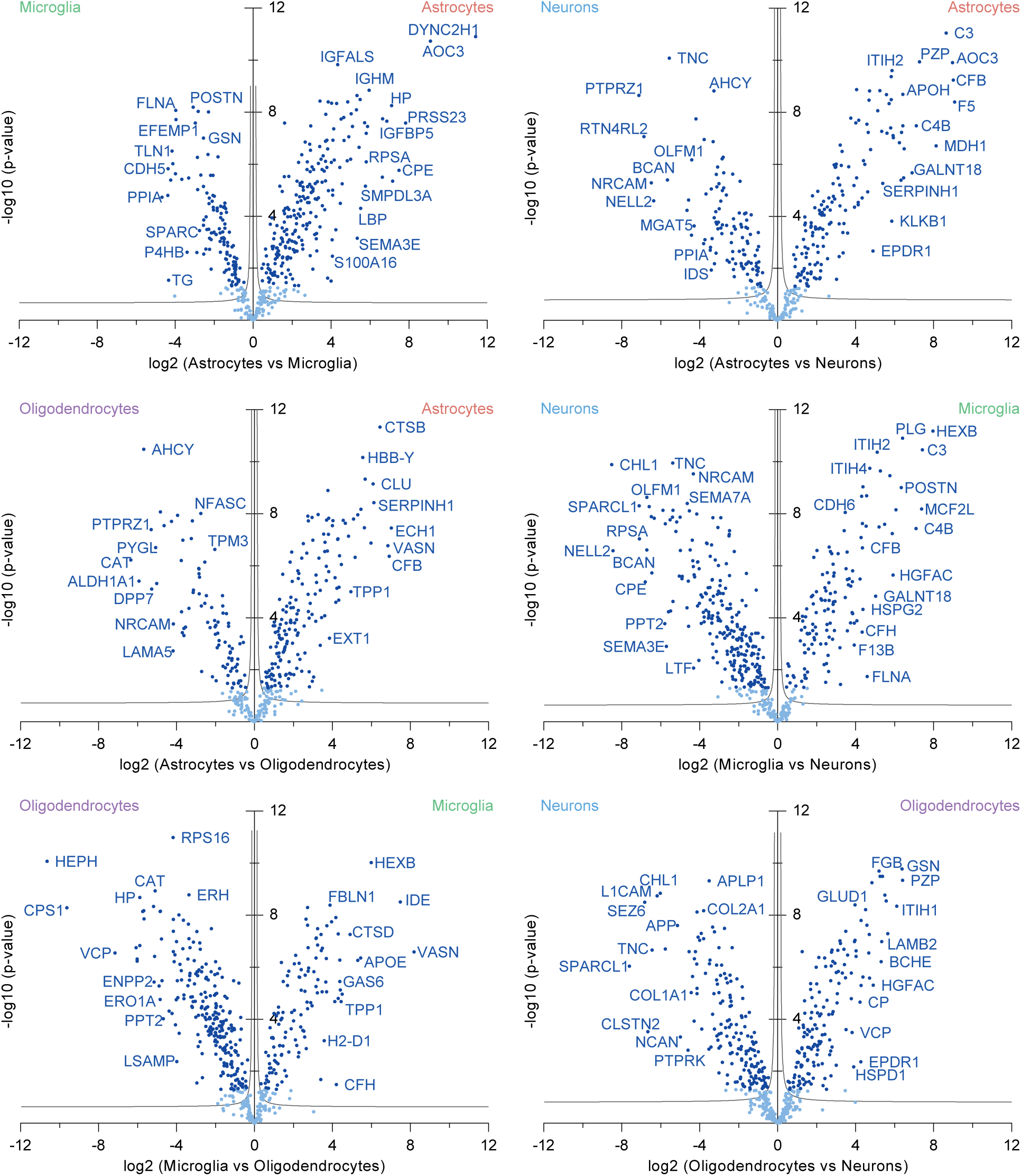
Diversity of cell type-resolved mouse brain glyco-secretome resource. Pairwise comparison of the secretomes of the four different cell types. Volcano plot indicating the proteins in the conditioned media of the different primary brain cells analyzed with the hiSPECS DIA method. The negative log10 transformed p-value of each protein is plotted against its log2 fold change comparing all investigated cell types with each other. Significantly regulated proteins (p-value< 0.05) are indicated in dark blue. The grey hyperbolic curves depict a permutation based false discovery rate estimation (p = 0.05; s0 = 0.1).

**Supplementary Fig. 4:**
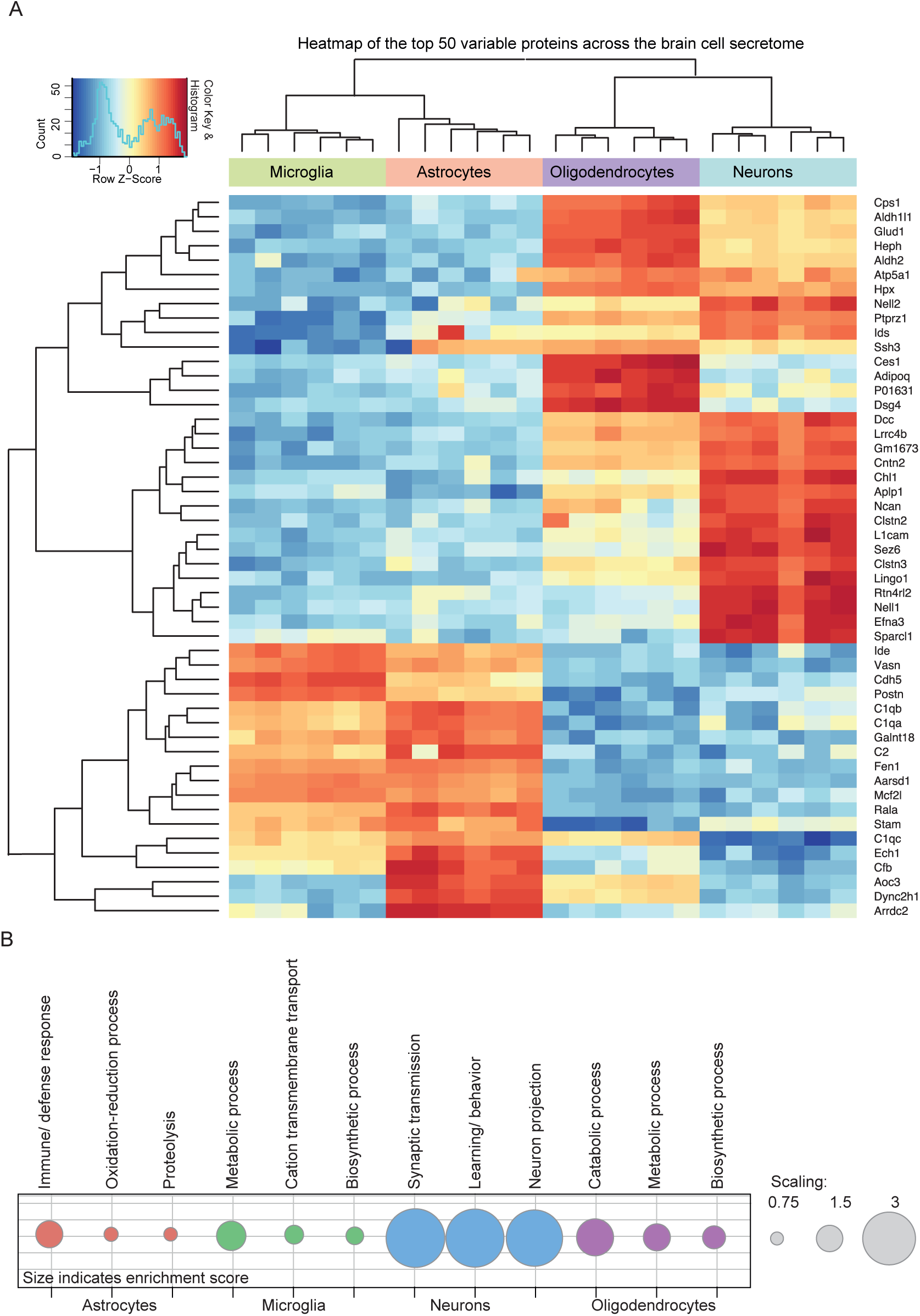
Top 50 cell type-specifically enriched proteins in the secretome resource. **A)** Heat map of the top 50 differentially secreted proteins (Bonferroni p.adj <0.05) across the four cell types from hierarchical clustering. For the missing protein quantification data, an imputation approach was undertaken using data missing at random within a left-shifted Gaussian distribution by 1.8 standard deviation. The rows represent the differentially secreted proteins and the columns represent the cell types with their replicates. The colors represent log-scaled protein levels with blue indicating the lowest, white indicating intermediate, and red indicating the highest protein levels. **B)** Functional annotation clustering with DAVID 6.8 (Huang da et al., 2009a, Huang da et al., 2009b) for gene ontology term biological process (FAT) of the cell type-specific secretome proteins (Supplementary Table 4). All proteins detected in the hiSPECS brain secretome study have been chosen as the background. The dot sizes indicate the enrichment score.

**Supplementary Fig. 5:**
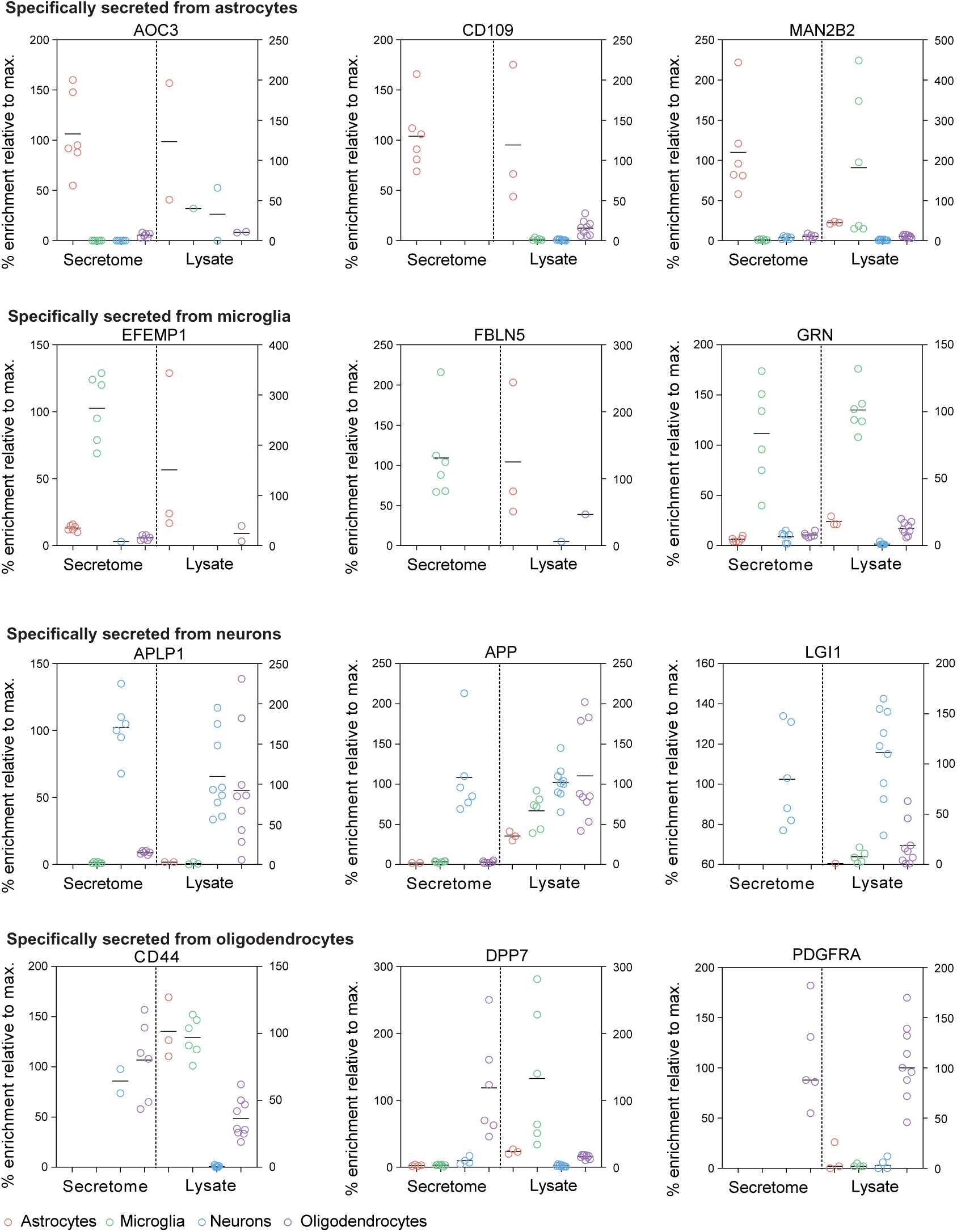
Protein levels in the brain cell secretome vs lysate proteome. Comparison of the iSPECS secretome resource and lysate data by (Sharma et al., 2015). The % enrichment is indicated normalized to the average of the most abundant cell type. For example, APLP1 is similarly abundant in lysates of neurons and oligodendrocytes, but only secreted to a relevant extent from neurons. Thus, APLP1 is classified as a cell type-specifically secreted protein.

**Supplementary Fig. 6:**
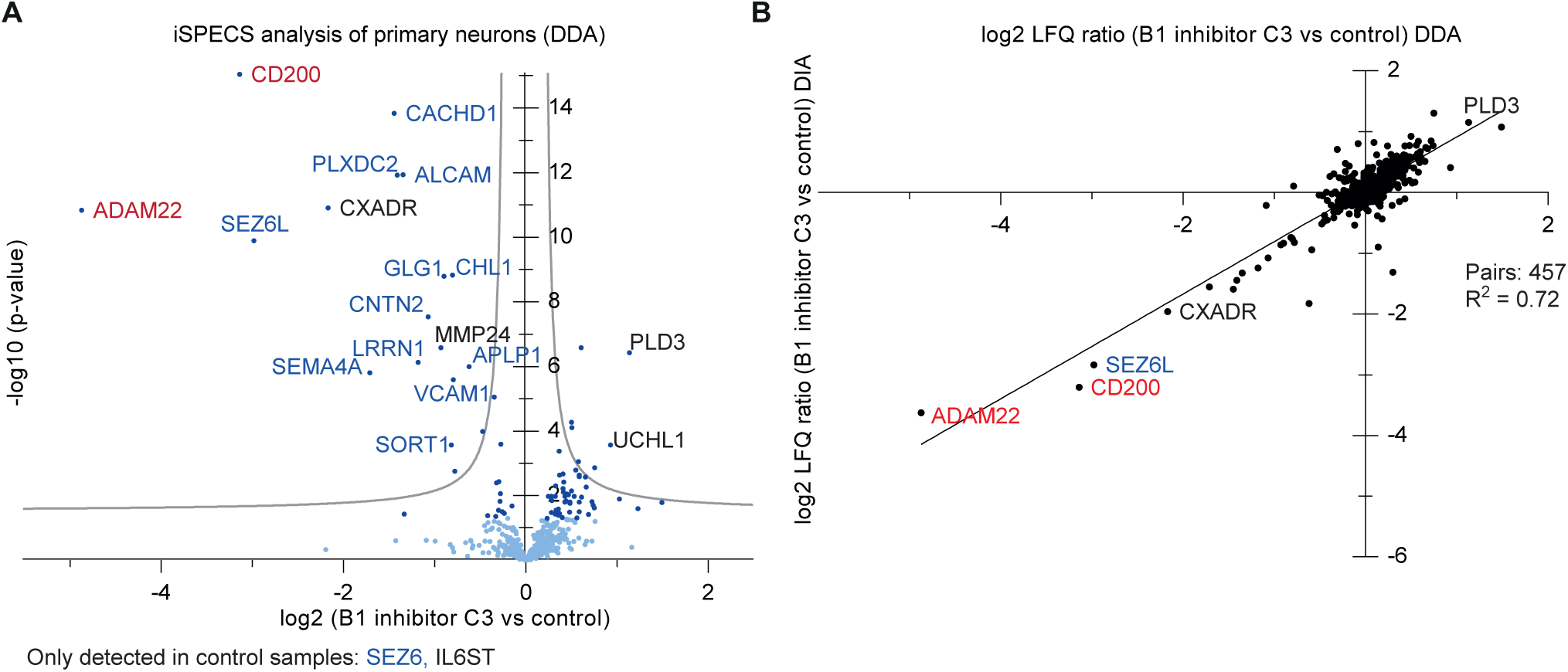
Substrate candidate identification of the Alzheimeŕs protease BACE1 as proof of principle of the iSPECS method. **A)** Same experiment as in Fig. 2A,B, but using DDA instead of DIA for mass spectrometric data acquisition. The volcano plot shows changes in protein levels in the secretome of primary cultured neurons upon BACE1 inhibitor C3 treatment using the iSPECS DDA method. The negative log10 transformed p-value of each protein is plotted against its log2 fold change comparing inhibitor treated and control condition. The grey hyperbolic curves depict a permutation based false discovery rate estimation (p = 0.05; s0 = 0.1). Significantly regulated proteins (p<0.05) are indicated with a dark blue dot and known BACE1 substrates are indicated with blue letters. The two newly validated BACE1 substrates CD200 and ADAM22 are indicated in red. SEZ6 and IL6ST are not depicted in the volcano plot because they were only identified in the control condition. **B)** Correlation of the hiSPECS DDA to DIA method. Plotted are the log2 fold changes between C3 treatment and DMSO control samples (N=11).

**Supplementary Fig. 7:**
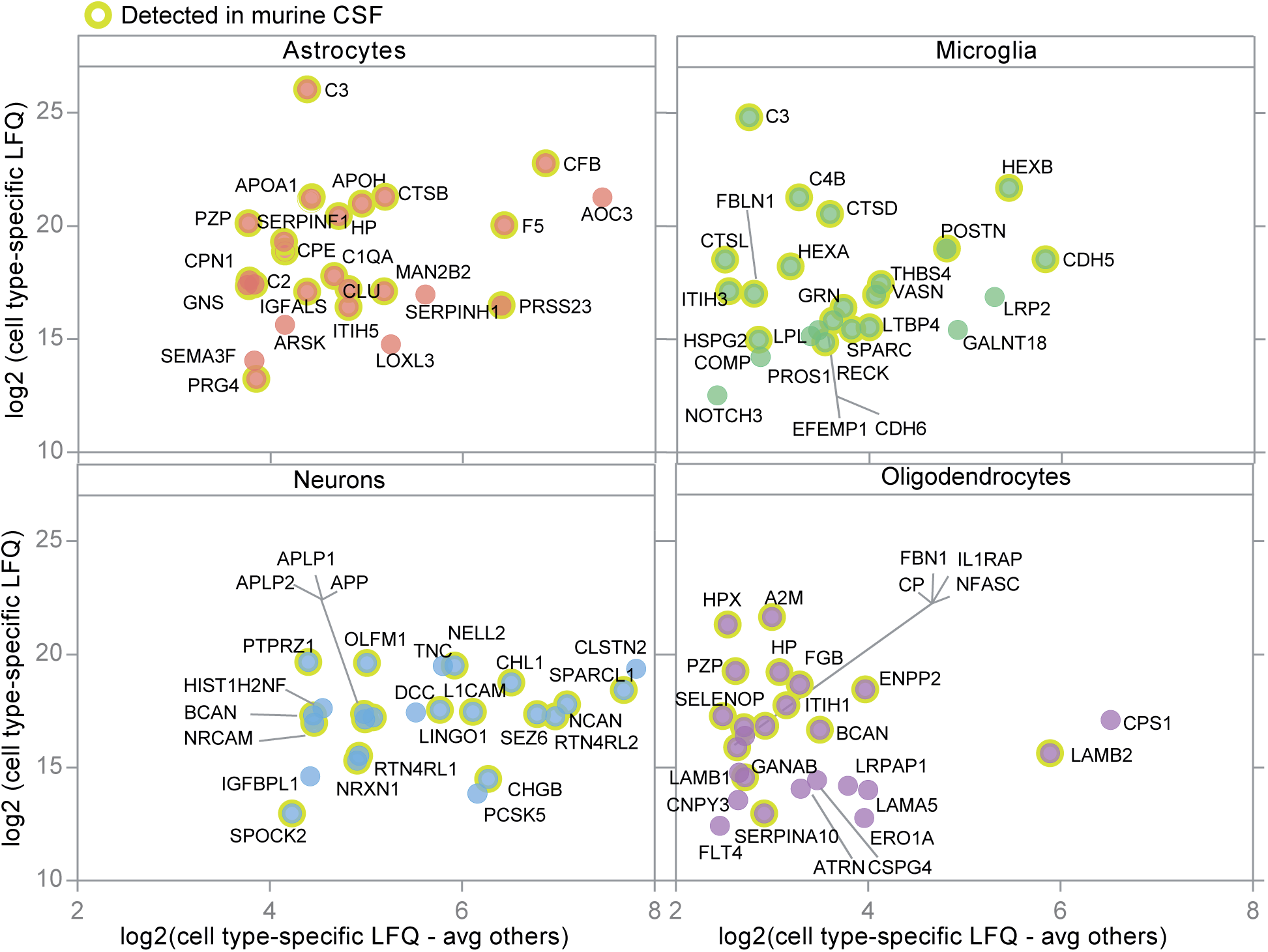
Top25 enriched proteins in the secretome of astrocytes, microglia, neurons and oligodendrocytes. Representation of the top 25 glycoproteins enriched in the secretome of one cell type. The average log2 LFQ intensities are plotted against the log2 LFQ ratios of the cell type specific-abundance subtracted from the average of the other cell types. Therefore, the values on the y-axis roughly indicate the abundance within a celĺs secretome, whereas the values on the x-axis show the enrichment compared to the other cell types in log2 scale. Yellow circles indicate proteins which are also detected in murine CSF. One example is HEXB in the microglia panel with an average LFQ value of 21.2 (y-axis) and a value of 5.4 on the x-axis. This indicates that the protein has a 2^5.4^-fold higher level in the secretome of microglia compared to the average of its levels in the secretome of the other three cell types.

**Table S7:** List of BACE1 substrate candidates. Summary of the significantly reduced proteins (p-values < 0.05) in the secretome of neurons upon pharmacological inihibtion of BACE1 with C3 identified by the hiSPECS DIA method (Figure 4b). Listed are the gene names, UniProt accession, ratio between C3 vs control samples (DMSO), p-value and the topology of the proteins. Other proteomic studies are highlighted which previously showed a reduction of the proteins upon BACE1 inhibition in primary neurons or murine CSF.

| Gene Name | UniProt Ac. | T-test p-value | Ratio (C3/Ctr) | Topology | Publication |
| --- | --- | --- | --- | --- | --- |
| Sez6 | Q7TSK2 | 3.04E-07 | 0.06 | TM1 |  |
| Adam22 | Q9R1V6 | 2.55E-15 | 0.08 | TM1 | Kuhn et al., 2012 |
| Cd200 | O54901 | 1.04E-21 | 0.11 | TM1 |  |
| Sez6l | Q6P1D5 | 4.25E-06 | 0.14 | TM1 | Kuhn et al., 2012 |
| Cxadr | P97792 | 2.83E-13 | 0.26 | TM1 |  |
| Il6st | Q00560 | 2.54E-03 | 0.27 | TM1 |  |
| Aplp1 | Q03157 | 1.60E-08 | 0.28 | TM1 | Kuhn et al., 2012; Zhou et al., 2012; Dislich et al., 2015 |
| Cachd1 | Q6PDJ1 | 1.46E-12 | 0.33 | TM1 | Kuhn et al., 2012 |
| Sema4a | Q62178 | 2.17E-07 | 0.34 | TM1 |  |
| Plxdc2 | Q9DC11 | 4.07E-12 | 0.37 | TM1 | Kuhn et al., 2012; Zhou et al., 2012; Dislich et al., 2015 |
| Alcam | Q61490 | 1.37E-05 | 0.40 | TM1 |  |
| Lrrn1 | Q61809 | 2.65E-12 | 0.42 | TM1 | Kuhn et al., 2012 |
| Cntn2 | Q61330 | 8.23E-14 | 0.47 | GPI | Kuhn et al., 2012; Zhou et al., 2012; Dislich et al., 2015 |
| Mmp24 | Q9R0S2 | 4.96E-05 | 0.55 | TM1 |  |
| Glg1 | Q61543 | 5.19E-10 | 0.56 | TM1 | Kuhn et al., 2012,9 |
| Fgfr1 | P16092 | 4.61E-05 | 0.57 | TM1 | Kuhn et al., 2012; Zhou et al., 2012; Dislich et al., 2015 |
| Vcam1 | P29533 | 1.13E-07 | 0.59 | TM1 |  |
| Chl1 | P70232 | 1.64E-09 | 0.60 | TM1 | Kuhn et al., 2012; Zhou et al., 2012; Dislich et al., 2015 |
| Sort1 | Q6PHU5 | 1.25E-04 | 0.60 | TM1 |  |
| Sema4b | Q62179 | 3.17E-07 | 0.70 | TM1 | Kuhn et al., 2012 |
| Col4a1 | P02463 | 2.50E-03 | 0.75 |  |  |
| Itm2b | O89051 | 1.77E-02 | 0.78 | TM2 |  |
| Met | P16056 | 7.93E-06 | 0.80 | TM1 |  |
| Col4a2 | P08122 | 1.72E-03 | 0.81 |  |  |
| Robo1 | O89026 | 4.41E-03 | 0.82 | TM1 |  |
| Adam12 | Q61824 | 4.85E-02 | 0.82 | TM1 |  |
| Tyro3 | P55144 | 5.02E-02 | 0.83 | TM1 |  |
| Tnfrsf21 | Q9EPU5 | 1.62E-02 | 0.84 | TM1 |  |
| Sema6c | Q9WTM3 | 7.94E-04 | 0.85 | TM1 |  |
| St6galnac5 | Q9QYJ1 | 4.80E-02 | 0.86 | TM2 |  |
| Mdga1 | Q0PMG2 | 2.57E-03 | 0.86 | GPI |  |
| Ptprg | Q05909 | 9.09E-04 | 0.88 | TM1 |  |
| Rgma | Q6PCX7 | 3.51E-02 | 0.91 | GPI |  |

## References

1. Bai B, Wang X, Li Y, Chen PC, Yu K, Dey KK, Yarbro JM, Han X, Lutz BM, Rao S, Jiao Y, Sifford JM, Han J, Wang M, Tan H, Shaw TI, Cho JH, Zhou S, Wang H, Niu M et al. (2020) Deep Multilayer Brain Proteomics Identifies Molecular Networks in Alzheimer’s Disease Progression. Neuron 105: 975–991.e7

2. Blank B, von Blume J (2017) Cab45-Unraveling key features of a novel secretory cargo sorter at the trans-Golgi network. European journal of cell biology 96: 383–390

3. Bruderer R, Bernhardt OM, Gandhi T, Miladinović SM, Cheng L-Y, Messner S, Ehrenberger T, Zanotelli V, Butscheid Y, Escher C, Vitek O, Rinner O, Reiter L (2015) Extending the limits of quantitative proteome profiling with data-independent acquisition and application to acetaminophen-treated three-dimensional liver microtissues. Molecular & cellular proteomics : MCP 14: 1400–1410

4. Chatr-Aryamontri A, Breitkreutz BJ, Oughtred R, Boucher L, Heinicke S, Chen D, Stark C, Breitkreutz A, Kolas N, O’Donnell L, Reguly T, Nixon J, Ramage L, Winter A, Sellam A, Chang C, Hirschman J, Theesfeld C, Rust J, Livstone MS et al. (2015) The BioGRID interaction database: 2015 update. Nucleic acids research 43: D470–8

5. Chitramuthu BP, Bennett HPJ, Bateman A (2017) Progranulin: a new avenue towards the understanding and treatment of neurodegenerative disease. Brain 140: 3081–3104

6. Dancourt J, Barlowe C (2010) Protein sorting receptors in the early secretory pathway. Annual review of biochemistry 79: 777–802

7. Daria A, Colombo A, Llovera G, Hampel H, Willem M, Liesz A, Haass C, Tahirovic S (2017) Young microglia restore amyloid plaque clearance of aged microglia. The EMBO journal 36: 583–603

8. Deshmukh AS, Cox J, Jensen LJ, Meissner F, Mann M (2015) Secretome Analysis of Lipid-Induced Insulin Resistance in Skeletal Muscle Cells by a Combined Experimental and Bioinformatics Workflow. Journal of proteome research 14: 4885–95

9. Diaz-Papkovich A, Anderson-Trocme L, Ben-Eghan C, Gravel S (2019) UMAP reveals cryptic population structure and phenotype heterogeneity in large genomic cohorts. PLoS genetics 15: e1008432

10. Dislich B, Wohlrab F, Bachhuber T, Muller SA, Kuhn PH, Hogl S, Meyer-Luehmann M, Lichtenthaler SF (2015) Label-free Quantitative Proteomics of Mouse Cerebrospinal Fluid Detects beta-Site APP Cleaving Enzyme (BACE1) Protease Substrates In Vivo. Molecular & cellular proteomics : MCP 14: 2550–63

11. Eichelbaum K, Winter M, Berriel Diaz M, Herzig S, Krijgsveld J (2012) Selective enrichment of newly synthesized proteins for quantitative secretome analysis. Nature biotechnology 30: 984–90

12. Ewers M, Franzmeier N, Suarez-Calvet M, Morenas-Rodriguez E, Caballero MAA, Kleinberger G, Piccio L, Cruchaga C, Deming Y, Dichgans M, Trojanowski JQ, Shaw LM, Weiner MW, Haass C (2019) Increased soluble TREM2 in cerebrospinal fluid is associated with reduced cognitive and clinical decline in Alzheimer’s disease. Science translational medicine 11

13. Fang P, Wang XJ, Xue Y, Liu MQ, Zeng WF, Zhang Y, Zhang L, Gao X, Yan GQ, Yao J, Shen HL, Yang PY (2016) In-depth mapping of the mouse brain N-glycoproteome reveals widespread N-glycosylation of diverse brain proteins. Oncotarget 7: 38796–38809

14. Gillet LC, Navarro P, Tate S, Rost H, Selevsek N, Reiter L, Bonner R, Aebersold R (2012) Targeted data extraction of the MS/MS spectra generated by data-independent acquisition: a new concept for consistent and accurate proteome analysis. Molecular & cellular proteomics : MCP 11: O111.016717

15. Gosselin D, Skola D, Coufal NG, Holtman IR, Schlachetzki JCM, Sajti E, Jaeger BN, O’Connor C, Fitzpatrick C, Pasillas MP, Pena M, Adair A, Gonda DD, Levy ML, Ransohoff RM, Gage FH, Glass CK (2017) An environment-dependent transcriptional network specifies human microglia identity. Science (New York, NY) 356

16. Huang da W, Sherman BT, Lempicki RA (2009a) Bioinformatics enrichment tools: paths toward the comprehensive functional analysis of large gene lists. Nucleic acids research 37: 1–13

17. Huang da W, Sherman BT, Lempicki RA (2009b) Systematic and integrative analysis of large gene lists using DAVID bioinformatics resources. Nature protocols 4: 44–57

18. Huang YA, Zhou B, Nabet AM, Wernig M, Sudhof TC (2019) Differential Signaling Mediated by ApoE2, ApoE3, and ApoE4 in Human Neurons Parallels Alzheimer’s Disease Risk. J Neurosci 39: 7408–7427

19. Johnson ECB, Dammer EB, Duong DM, Ping L, Zhou M, Yin L, Higginbotham LA, Guajardo A, White B, Troncoso JC, Thambisetty M, Montine TJ, Lee EB, Trojanowski JQ, Beach TG, Reiman EM, Haroutunian V, Wang M, Schadt E, Zhang B et al. (2020) Large-scale proteomic analysis of Alzheimer’s disease brain and cerebrospinal fluid reveals early changes in energy metabolism associated with microglia and astrocyte activation. Nature Medicine

20. Joshi HJ, Jorgensen A, Schjoldager KT, Halim A, Dworkin LA, Steentoft C, Wandall HH, Clausen H, Vakhrushev SY (2018) GlycoDomainViewer: a bioinformatics tool for contextual exploration of glycoproteomes. Glycobiology 28: 131–136

21. Kammers K, Cole RN, Tiengwe C, Ruczinski I (2015) Detecting Significant Changes in Protein Abundance. EuPA open proteomics 7: 11–19

22. Kleifeld O, Doucet A, auf dem Keller U, Prudova A, Schilling O, Kainthan RK, Starr AE, Foster LJ, Kizhakkedathu JN, Overall CM (2010) Isotopic labeling of terminal amines in complex samples identifies protein N-termini and protease cleavage products. Nature biotechnology 28: 281–8

23. Kleifeld O, Doucet A, Prudova A, auf dem Keller U, Gioia M, Kizhakkedathu JN, Overall CM (2011) Identifying and quantifying proteolytic events and the natural N terminome by terminal amine isotopic labeling of substrates. Nature protocols 6: 1578–611

24. Kuhn PH, Colombo AV, Schusser B, Dreymueller D, Wetzel S, Schepers U, Herber J, Ludwig A, Kremmer E, Montag D, Muller U, Schweizer M, Saftig P, Brase S, Lichtenthaler SF (2016) Systematic substrate identification indicates a central role for the metalloprotease ADAM10 in axon targeting and synapse function. eLife 5

25. Kuhn PH, Koroniak K, Hogl S, Colombo A, Zeitschel U, Willem M, Volbracht C, Schepers U, Imhof A, Hoffmeister A, Haass C, Rossner S, Brase S, Lichtenthaler SF (2012) Secretome protein enrichment identifies physiological BACE1 protease substrates in neurons. The EMBO journal 31: 3157–68

26. Lee TH, Cheng KK, Hoo RL, Siu PM, Yau SY (2019) The Novel Perspectives of Adipokines on Brain Health. International journal of molecular sciences 20

27. Lichtenthaler SF (2012) Cell biology. Sheddase gets guidance. Science (New York, NY 335: 179–80

28. Lichtenthaler SF, Lemberg MK, Fluhrer R (2018) Proteolytic ectodomain shedding of membrane proteins in mammals-hardware, concepts, and recent developments. The EMBO journal 37

29. Liu MQ, Zeng WF, Fang P, Cao WQ, Liu C, Yan GQ, Zhang Y, Peng C, Wu JQ, Zhang XJ, Tu HJ, Chi H, Sun RX, Cao Y, Dong MQ, Jiang BY, Huang JM, Shen HL, Wong CCL, He SM et al. (2017) pGlyco 2.0 enables precision N-glycoproteomics with comprehensive quality control and one-step mass spectrometry for intact glycopeptide identification. Nature communications 8: 438

30. Ludwig C, Gillet L, Rosenberger G, Amon S, Collins BC, Aebersold R (2018) Data-independent acquisition-based SWATH-MS for quantitative proteomics: a tutorial. Molecular systems biology 14: e8126

31. Meissner F, Scheltema RA, Mollenkopf HJ, Mann M (2013) Direct proteomic quantification of the secretome of activated immune cells. Science (New York, NY) 340: 475–8

32. O’Brien RJ, Wong PC (2011) Amyloid precursor protein processing and Alzheimer’s disease. Annu Rev Neurosci 34: 185–204

33. Olsson B, Lautner R, Andreasson U, Ohrfelt A, Portelius E, Bjerke M, Holtta M, Rosen C, Olsson C, Strobel G, Wu E, Dakin K, Petzold M, Blennow K, Zetterberg H (2016) CSF and blood biomarkers for the diagnosis of Alzheimer’s disease: a systematic review and meta-analysis. The Lancet Neurology 15: 673–684

34. Pigoni M, Wanngren J, Kuhn PH, Munro KM, Gunnersen JM, Takeshima H, Feederle R, Voytyuk I, De Strooper B, Levasseur MD, Hrupka BJ, Muller SA, Lichtenthaler SF (2016) Seizure protein 6 and its homolog seizure 6-like protein are physiological substrates of BACE1 in neurons. Molecular neurodegeneration 11: 67

35. Pinero J, Bravo A, Queralt-Rosinach N, Gutierrez-Sacristan A, Deu-Pons J, Centeno E, Garcia-Garcia J, Sanz F, Furlong LI (2017) DisGeNET: a comprehensive platform integrating information on human disease-associated genes and variants. Nucleic acids research 45: D833–d839

36. Rappsilber J, Ishihama Y, Mann M (2003) Stop and go extraction tips for matrix-assisted laser desorption/ionization, nanoelectrospray, and LC/MS sample pretreatment in proteomics. Analytical chemistry 75: 663–70

37. Ritchie ME, Phipson B, Wu D, Hu Y, Law CW, Shi W, Smyth GK (2015) limma powers differential expression analyses for RNA-sequencing and microarray studies. Nucleic acids research 43: e47

38. Schindler SE, Li Y, Todd KW, Herries EM, Henson RL, Gray JD, Wang G, Graham DL, Shaw LM, Trojanowski JQ, Hassenstab JJ, Benzinger TLS, Cruchaga C, Jucker M, Levin J, Chhatwal JP, Noble JM, Ringman JM, Graff-Radford NR, Holtzman DM et al. (2019) Emerging cerebrospinal fluid biomarkers in autosomal dominant Alzheimer’s disease. Alzheimer’s & dementia : the journal of the Alzheimer’s Association 15: 655–665

39. Schira-Heinen J, Grube L, Waldera-Lupa DM, Baberg F, Langini M, Etemad-Parishanzadeh O, Poschmann G, Stuhler K (2019) Pitfalls and opportunities in the characterization of unconventionally secreted proteins by secretome analysis. Biochimica et biophysica acta Proteins and proteomics 1867: 140237

40. Schlage P, Kockmann T, Kizhakkedathu JN, auf dem Keller U (2015) Monitoring matrix metalloproteinase activity at the epidermal-dermal interface by SILAC-iTRAQ-TAILS. Proteomics 15: 2491–502

41. Sharma K, Schmitt S, Bergner CG, Tyanova S, Kannaiyan N, Manrique-Hoyos N, Kongi K, Cantuti L, Hanisch UK, Philips MA, Rossner MJ, Mann M, Simons M (2015) Cell type- and brain region-resolved mouse brain proteome. Nature neuroscience 18: 1819–31

42. Stachel SJ, Coburn CA, Steele TG, Jones KG, Loutzenhiser EF, Gregro AR, Rajapakse HA, Lai MT, Crouthamel MC, Xu M, Tugusheva K, Lineberger JE, Pietrak BL, Espeseth AS, Shi XP, Chen-Dodson E, Holloway MK, Munshi S, Simon AJ, Kuo L et al. (2004) Structure-based design of potent and selective cell-permeable inhibitors of human beta-secretase (BACE-1). Journal of medicinal chemistry 47: 6447–50

43. Stiess M, Wegehingel S, Nguyen C, Nickel W, Bradke F, Cambridge SB (2015) A Dual SILAC Proteomic Labeling Strategy for Quantifying Constitutive and Cell-Cell Induced Protein Secretion. Journal of proteome research 14: 3229–3238

44. Stutzer I, Selevsek N, Esterhazy D, Schmidt A, Aebersold R, Stoffel M (2013) Systematic proteomic analysis identifies beta-site amyloid precursor protein cleaving enzyme 2 and 1 (BACE2 and BACE1) substrates in pancreatic beta-cells. The Journal of biological chemistry 288: 10536–47

45. Suarez-Calvet M, Araque Caballero MA, Kleinberger G, Bateman RJ, Fagan AM, Morris JC, Levin J, Danek A, Ewers M, Haass C, Dominantly Inherited Alzheimer N (2016) Early changes in CSF sTREM2 in dominantly inherited Alzheimer’s disease occur after amyloid deposition and neuronal injury. Sci Transl Med 8: 369ra178

46. UniProt Consortium T (2018) UniProt: the universal protein knowledgebase. Nucleic acids research 46: 2699

47. Vassar R, Bennett BD, Babu-Khan S, Kahn S, Mendiaz EA, Denis P, Teplow DB, Ross S, Amarante P, Loeloff R, Luo Y, Fisher S, Fuller J, Edenson S, Lile J, Jarosinski MA, Biere AL, Curran E, Burgess T, Louis JC et al. (1999) Beta-secretase cleavage of Alzheimer’s amyloid precursor protein by the transmembrane aspartic protease BACE. Science (New York, NY 286: 735–41.

48. Voytyuk I, Mueller SA, Herber J, Snellinx A, Moechars D, van Loo G, Lichtenthaler SF, De Strooper B (2018) BACE2 distribution in major brain cell types and identification of novel substrates. Life Sci Alliance 1: e201800026–e201800026

49. Wiita AP, Seaman JE, Wells JA (2014) Global analysis of cellular proteolysis by selective enzymatic labeling of protein N-termini. Methods in enzymology 544: 327–58

50. Wolfe CM, Fitz NF, Nam KN, Lefterov I, Koldamova R (2018) The Role of APOE and TREM2 in Alzheimer’s Disease-Current Understanding and Perspectives. International journal of molecular sciences 20: 81

51. Yan R, Bienkowski MJ, Shuck ME, Miao H, Tory MC, Pauley AM, Brashier JR, Stratman NC, Mathews WR, Buhl AE, Carter DB, Tomasselli AG, Parodi LA, Heinrikson RL, Gurney ME (1999) Membrane-anchored aspartyl protease with Alzheimer’s disease beta-secretase activity. Nature 402: 533–7

52. Yi MH, Zhang E, Kim JJ, Baek H, Shin N, Kim S, Kim SR, Kim HR, Lee SJ, Park JB, Kim Y, Kwon OY, Lee YH, Oh SH, Kim DW (2016) CD200R/Foxp3-mediated signalling regulates microglial activation. Scientific reports 6: 34901

53. Yin RH, Yu JT, Tan L (2015) The Role of SORL1 in Alzheimer’s Disease. Molecular neurobiology 51: 909–18

54. Zetterberg H, Bendlin BB (2020) Biomarkers for Alzheimer’s disease-preparing for a new era of disease-modifying therapies. Molecular psychiatry

55. Zhang Y, Chen K, Sloan SA, Bennett ML, Scholze AR, O’Keeffe S, Phatnani HP, Guarnieri P, Caneda C, Ruderisch N, Deng S, Liddelow SA, Zhang C, Daneman R, Maniatis T, Barres BA, Wu JQ (2014) An RNA-sequencing transcriptome and splicing database of glia, neurons, and vascular cells of the cerebral cortex. J Neurosci 34: 11929–11947

56. Zielinska DF, Gnad F, Wisniewski JR, Mann M (2010) Precision mapping of an in vivo N-glycoproteome reveals rigid topological and sequence constraints. Cell 141: 897–907

